# Determining the age of single cells using scMLEAge

**DOI:** 10.64898/2025.12.04.692166

**Authors:** Chanyue Hu, Matteo Pellegrini

## Abstract

Aging is a complex biological process marked by a gradual decline in physiological function that contributes to increased vulnerability to disease and mortality. Numerous studies have investigated the cellular and molecular aspects of aging at single-cell resolution, yet the heterogeneity of cellular aging in an individual remains poorly understood. To enhance our ability to study aging at the single cell level, we developed a statistical framework to predict the age of individual cells based on their transcriptomic profiles. Our Bayesian approach estimates the most likely age of a cell given its read counts. We applied the model to data from Tabula Muris Senis and examined organ- and cell-type-specific transcriptomic signatures of aging. Compared with standard regression-based methods, our framework achieved higher predictive accuracy. We show that *scMLEAge* is a powerful tool for dissecting the cellular heterogeneity of aging and age-related functional decline.

## 2 Introduction

Aging is a fundamental biological process that involves gradual decline in cellular and physiological function, ultimately leading to increased susceptibility to age-related diseases and mortality. To study this process, multiple molecular markers have been developed that track the underlying mechanisms of aging. In the last 20 years, one of the most influential approaches has been the use of DNA methylation data to construct multi-tissue epigenetic clocks [1–5]. These methylation clocks accurately predict the age of an individual from their methylation profiles.

In parallel to epigenetic clocks, transcriptomic approaches using bulk RNA sequencing have offered complementary insights into the aging process by identifying gene expression patterns that correlate with chronological age. Some of these clocks use linear regression to model age associated transcriptomic changes while others incorporate nonlinear methods (e.g. *BayesAge 2*.*0* [6]). Despite their utility, these approaches rely on pooling all the RNA signals across heterogeneous cell populations, which can obscure important cell type specific aging differences. This limitation is particularly significant as studies have shown that aging exhibits unique and asynchronous dynamics across tissues and cell types[7, 8].

The advent of single-cell RNA sequencing (scRNA-seq), a technology for profiling the transcriptome of individual cells, provides insights that were previously obscured by traditional bulk RNA sequencing methods. Several recent studies have applied scRNA-seq to aging research, identifying changes in cell type composition, shifts in gene expression, and age-associated markers across tissues and lifespan stages [9–11]. These advances have spurred the development of scRNA-based aging clocks capable of predicting the age of individual cells, or small collections of similar cells. These approaches assume that the age of cell is not necessarily the same as the age of the individual from which it is collected. In other words, within an individual, cells can have a variety of ages, reflecting an underlying aging trajectory of individual cells. However, the age a cell is not known ahead of time, and therefore these methods assume that a single cell aging clock can be trained by initially assuming that the age of a cell is that of the individual from which it is collected, and then using these models to predict the age of a single cell which could be different from that of the individual.

Generating single cell aging clocks is a challenging problem as the gene counts for single cells are usually much lower than in bulk data, and many genes are not observed at all. Moreover, the transcriptional state of a single cell is not known a priori, as it can deviate from the chronological age of the host organism. Buckley et al.[12], developed an aging clock that can accurately predict the age of a cell using the pseudo-bulk of cells in scRNA-seq expression. Other scRNA aging clocks have also been developed to predict the age of individual cells [13–15]. While these models mark an important step toward cell-level age prediction, they exhibit certain limitations across tissues and age ranges. The model by Zakar et al. employs an ElasticNet regression framework, which, although effective for feature selection, requires transformed, continuous expression values. This approach does not take into account the discrete nature of raw single-cell count data that are inherently sparse and zero-inflated. Zhu et al.’s clock is tailored to human supercentenarians, focusing on extremely high ages, therefore has limited applicability to broader or more diverse age ranges. Li et al. focuses on immune cells for age prediction in a study of samples from COVID patients, providing valuable insights into immunology but limiting the model’s generalizability across other tissues and systemic aging processes.

Here we present *scMLEAge*, a general statistical framework for predicting the age of individual cells from single-cell transcriptomic data across multiple tissues and age groups. Our approach estimates the maximum likelihood of the age of each cell based on the assumption that read counts can be modeled using a Poisson function, which is well-suited for count-based scRNA-seq data and accounts for inherent sparsity and variability of single cell data. We perform feature selection by tuning the number of genes included in each model, using a hyperparameter search strategy for model performance optimization. To provide an example of the utility of our method, we apply *scMLEAge* to the Tabula Muris Senis (TMS) dataset[9]. This single-cell transcriptomic atlas profiles over 350,000 cells covering 23 organs and 6 distinct age groups ranging from 1 to 30 months. Using *scMLEAge* we uncovered robust tissue-specific and cell-type-expression signatures associated with aging and demonstrate the ability to accurately estimate age at the single-cell level. Our framework provides a novel approach for dissecting transcriptional aging patterns with improved resolution, flexibility, and interpretability, advancing the toolkit for aging research in single-cell genomics.

## 3 Methods

### 3.1 Data Availability

We utilized the Tabula Muris Senis (TMS) dataset, a publicly available mouse single-cell RNA-seq dataset [9]. We selected bladder, bone-marrow, brain myeloid, brain non-myeloid, heart, kidney, limb-muscle, liver, and lung cells. The single-cell RNA sequencing data was obtained from the figshare link the Tabula Muris Consortium provided, https://doi.org/10.6084/m9.figshare.8273102.v2. The Tabula Muris atlas were generated using two different techniques: microtitre well plates (fluorescence-activated cell sorting; FACS) and microfluidic droplets. To avoid batch effects, we only used the droplet data. Since there is an inconsistency in ages between male and female samples in the data, only cells from 10 males are used in our models to remove sex effects. To ensure sufficient variability and data abundance for model training, we included only organs that contained at least three distinct age groups spanning from young (1–3 months) to old (24–30 months), and with a minimum of 100 cells per age group within each cell type. Furthermore, we removed cells in a cell type when a single age class comprised more than 80% of the total cell counts. This filtering ensured a more balanced representation of age groups across the dataset.

### 3.2 Constructing the Aging Clock

Our goal is to predict the age of a cell based on raw counts of individual cells. Each aging clock is constructed using a model tailored to each cell type within the organ of interest. Cells are split into training and testing sets using 5-fold cross-validation for each donor.

To estimate the frequency of each gene in each cell type in each organ, we first aggregate all cells associated with each category. For each cell type, we computed the frequency matrix *F* ∈ ℝ^*G×A*^ where *G* denotes the number of genes and *A* is the number of discrete age groups found in the training data. We fit the frequency to a linear regression model, and then we used the values of the model as the expected frequency 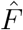. For feature selection, we ranked genes based on the Pearson correlation coefficient between normalized gene expression counts and chronological ages using the pearsonr method from the SciPy statistics package [16].

We used these expected gene frequencies for each age group to then estimate the age of individual cells. To do this, we first calculated the expected counts under the Poisson model for each gene *g* and age group *a* as:

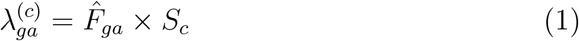

where the total expression count *S*_*c*_ is a function of *X*_*gc*_, which denotes the count of gene *g* in cell *c* and for a given cell *c*,:

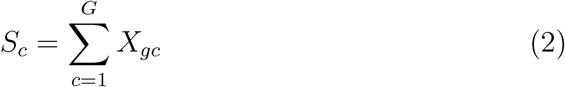

We can then compute the probability of observing the counts for each gene using a Poisson model:

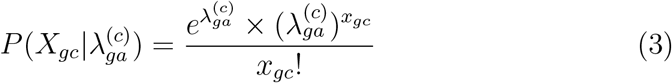

Finally, we take the maximum log-likelihood of our model as the predicted age group *â*_*c*_:

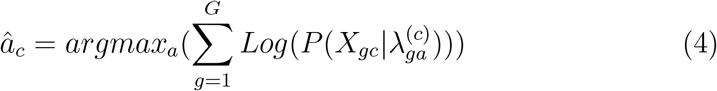

To optimize our hyperparameter corresponding to the number of genes to include in the model, we repeat this procedure using a power-of-two grid search over the list of the most correlated genes with age. Let 𝒢_*k*_ be the top 2^*k*^ genes sorted by correlation values with age. For each subset 𝒢_*k*_, we repeat this training and testing process and compute the squared Pearson correlation *r*^2^ by:

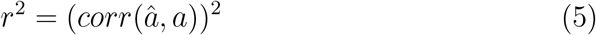

where

*â*: predicted age

*a*: true chronological age

The optimal number of genes was selected based on the value of *k* that maximized *r*^2^.

### 3.3 Age associated genes across cell types

We used two approaches to identify age associated genes across cell types. In the first approach to identify age-associated markers across multiple tissues (bladder, bone marrow, brain myeloid, brain non-myeloid, heart, kidney, limb muscle, liver, and lung) in the single-cell level, we computed Pearson’s correlation coefficients between the predicted ages of the cells and the expression levels of all genes in the TMS scRNA-seq dataset. For each gene, *p*-values were obtained and adjusted for multiple testing using the Benjamini–Hochberg procedure as implemented in the Python statsmodels package[17], applied across all tissues. The adjusted *p*-values were then log-transformed. Finally, genes were ranked by the sum of their log-transformed adjusted *p*-values across tissues, and the top 100 ranked genes were selected.

In a second approach we performed differential expression analysis (Wilcoxon rank-sum test, FDR ¡ 0.05) between cells of different predicted ages using scanpy [18] for each cell type within each tissue organ. Significant marker genes were identified using p-adjusted-value threshold of *<* 0.05. For each analysis, the top 20 genes were selected and combined across all of the cell types and tissues to assess preservation of markers between cell types and organs. Genes were ranked by *log*_2_ fold change values, with *p*-values used as a secondary sorting criterion.

## 4 Results

### 4.1 Building the scRNA clocks: *scMLEAge* framework

We developed single cell aging clocks in multiple tissues in the TMS data, including bladder, bone marrow, brain, heart, kidney, limb-muscle, liver and lung (Figure 1a). We only selected cell types with a balanced population of cells across age groups in order to have a sufficient numbers of cells across different age groups (see Methods). For each organ, we developed cell-type-specific aging clocks, using 5-fold cross-validation stratified by donor. Each model optimizes the number of genes to include. After selecting the best-performing model from the training data, we tested our model’s accuracy on the testing data. Overall, our models achieved Pearson’s R squared values ranging from 0.17 to 0.95 (Figure 1b&S Figure 1-9). Notably, the majority of models included 8192 genes, approximately half the total number of observed genes (Figure 1c), although some models used much smaller numbers of genes.

**Figure 1:**
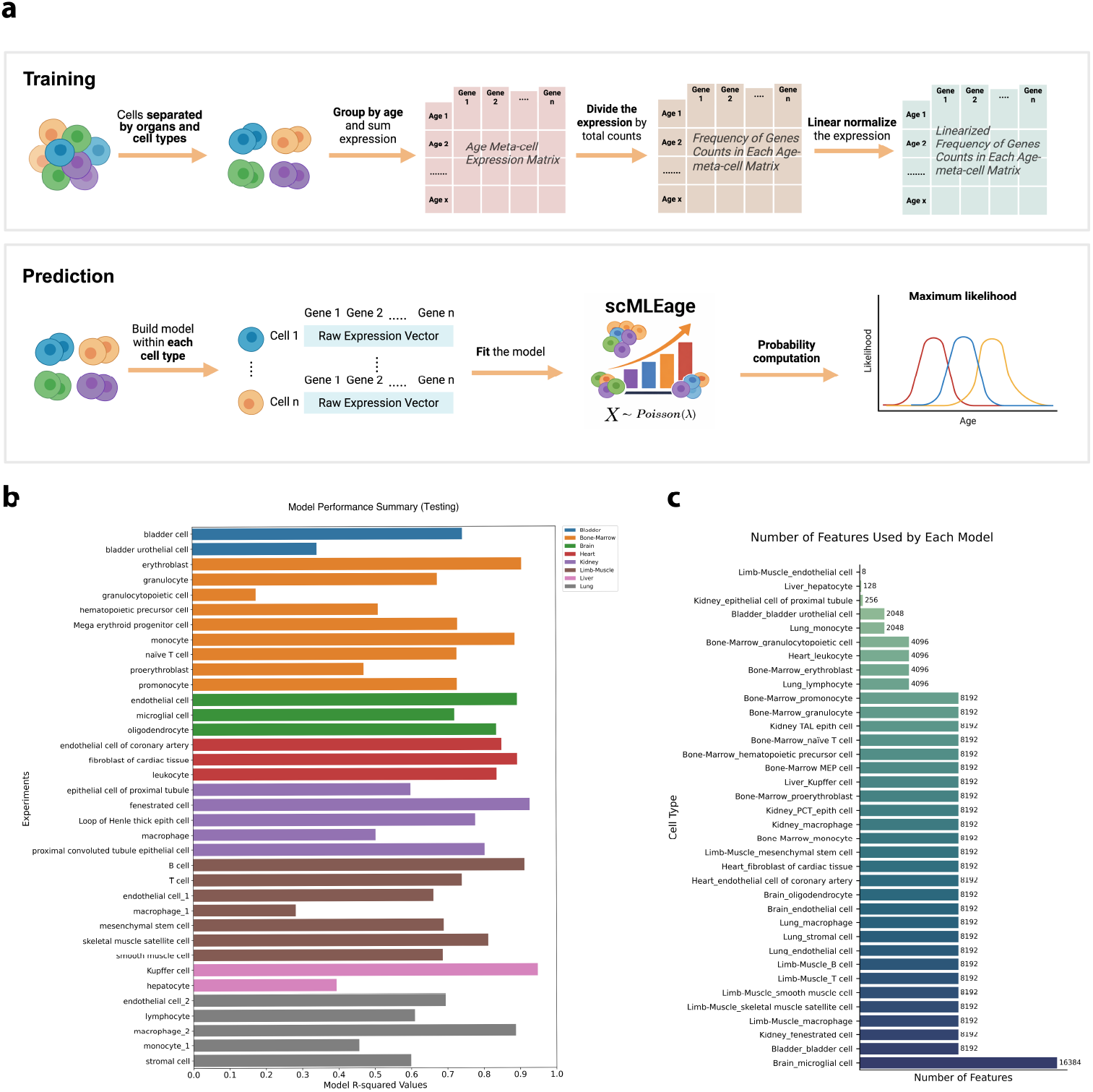
*scMLEAge* aging clocks. **a**: Schematic of the model. Training and prediction are cell-type-specific. During training, gene expression is aggregated by age group, normalized, and transformed into gene frequency distributions, which are then used to build the model. During prediction, the raw count vector of a single cell is used as the input to a Poisson model to estimate the probability distribution over ages, and selecting the age that maximizes the likelihood. **b**: Performance of individual models across cell types on test samples. **c**: The number of features (genes) used in each model.

#### 4.1.1 Method Comparison with ElasticNet and Lasso Model

To benchmark our approach against a widely used method in the field, we implemented an ElasticNet and Lasso cross validation model on limb muscle, kidney and lung cells from TMS. The expression counts were normalized before training and testing. The ElasticNet model used the following parameters: l1 ratio = 0.5, cv = 5, and max iter = 500. Alphas are specified as none (alphas = None) in order for the model to use cross validation to internally tune for the best alpha value. For the Lasso model, the alpha was also specified as none for the model itself to determine the best alpha value. 5-fold cross-validation stratified by donor was used to evaluate performance. Model accuracy was assessed by the squared Pearson correlation (*R*^2^). Compared with ElasticNet and Lasso, *scMLEAge* achieved superior performance across most cell types, including T cells from limb-muscle, B cells from limb-muscle, fenestrated cells from kidney, epithelial cells of proximal tubule from kidney, endothelial cells from lung, stromal cells and lymphocytes from lung. (Figure 2).

**Figure 2:**
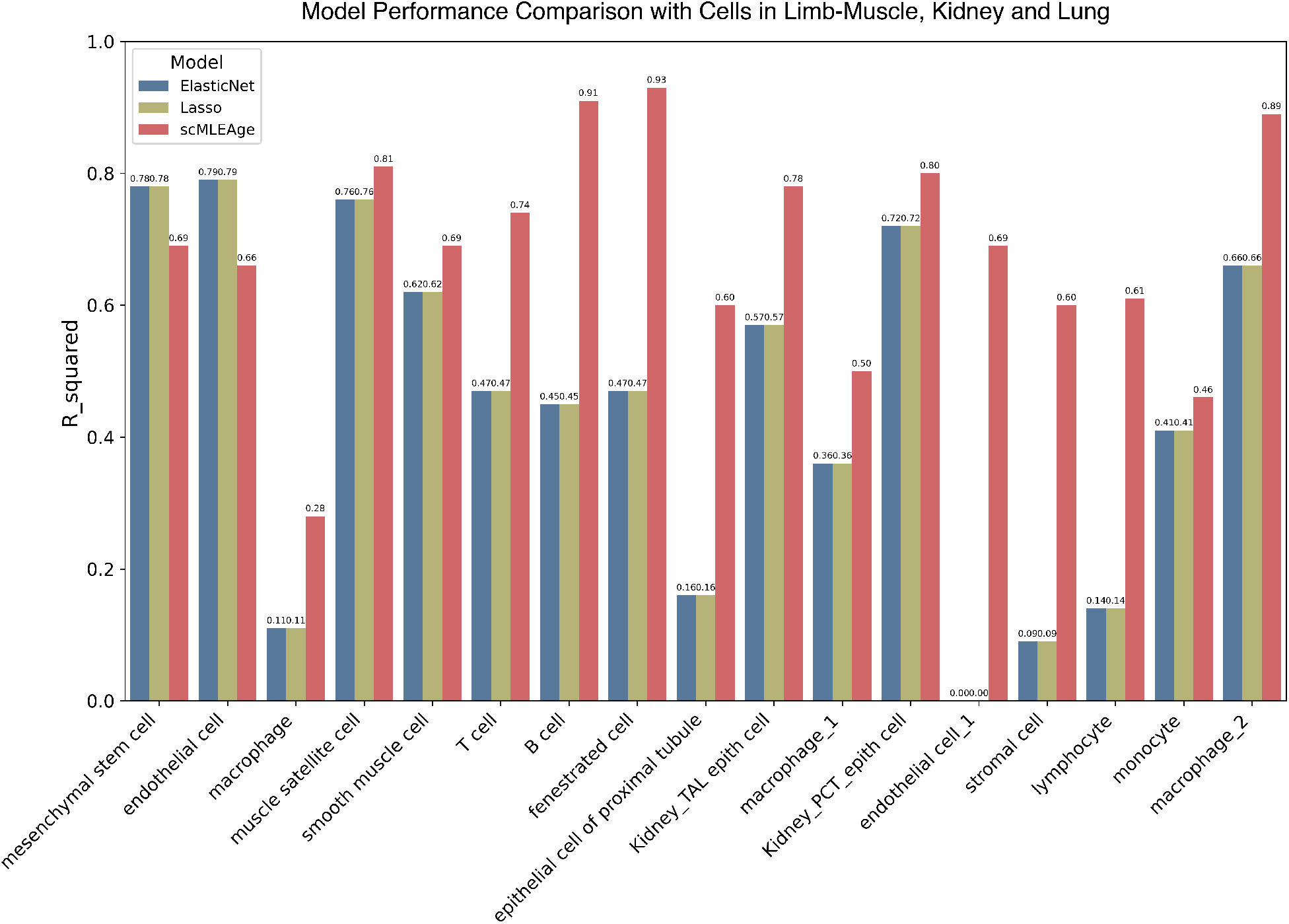
scMLEAge and ElasticNet models applied to TMS cells in Limb-muscle, Kidney and Lung. Bar plot showing the R squared values for each cell-type-specific model using scMLEAge and ElasticNet

#### 4.1.2 Limb-Muscle Skeletal Satellite Cells

As a first example of the cell type specific aging models we constructed using *scMLEAge*, we show the results for limb muscle skeletal satellite cells (LMuSSs). Using 11 mice of 1 month, 18 months, 24 months and 30 months, we achieved an R-squared value of 0.81 during training and 0.81 during testing (Figure 3a). Skeletal satellite cells in limb muscles are a population of myogenic stem cells responsible for regenerating skeletal muscle tissue, residing within the muscle fibers of the limbs and their population decreases with aging[19, 20].

**Figure 3:**
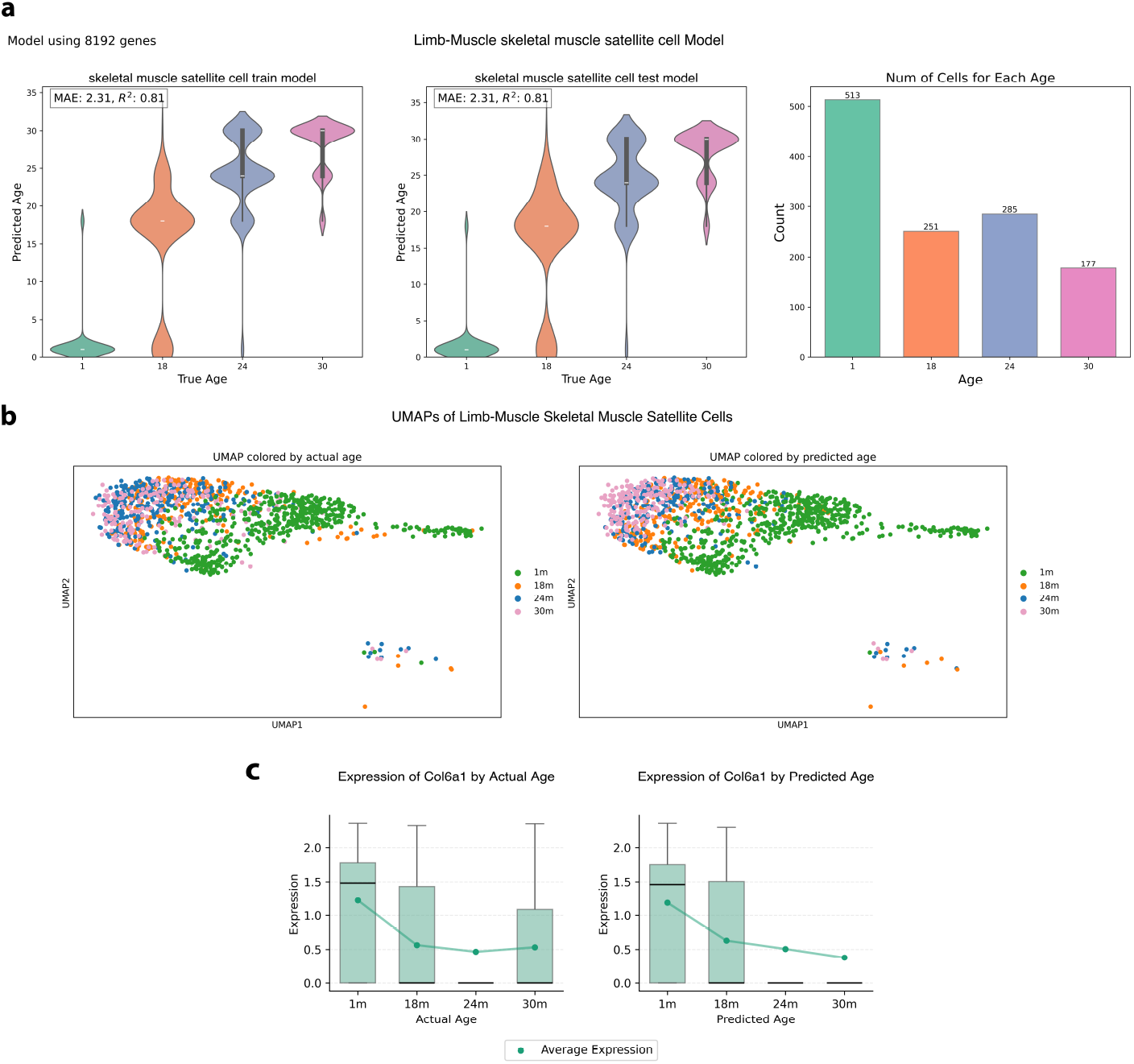
TMS Limb-muscle satellite cell Model. **a**: Violin plots of true age versus predicted age for limb-muscle satellite cell training data (left) and testing data (middle), along with a bar plot showing the number of cells at each age in training and test sets (right). **b**: UMAPs colored by actual ages (left) and predicted age (right). **c**: Box plots of COL6A1 expression across actual ages (left) and predicted ages (right), with dotted trend lines indicating average expression change with ages.

From the Uniform Manifold Approximation and Projection (UMAP) of the cells, we were able to visualize the heterogeneity of cells with respect to both actual ages and predicted ages. Cells follow a clear trajectory from the age of 1 month to 18 months. While the 24-month and 30-month cells are mixed together in the UMAP of actual ages (Figure 3b), in the UMAP of the predicted ages (Figure 3b), cells from 24 and 30-month-old mice do not fully overlap but instead distribute along a gradient. This suggests that our approach is able to assign ages that better separate along an inherent transcriptional age associated gradient.

We examined the top 100 most age-correlated genes to predicted ages and cross-referenced them with literature studies in mammal agings. One of the most highly correlated genes is COL6A1, a collagen VI gene whose protein product is secreted to the extracellular matrix, which is ranked at 78th on our list. Pearson correlation analysis revealed a moderate negative association with age (*r* = −0.40). *COL6A1* expression and average expression in each age group declined with both actual and predicted ages (Fig. 3c), with only the predicted-age axis displaying a monotonic trajectory of decline (Fig. 3c). Additionally, we observed 3 other collagen genes from the top 100 list (Fig. S1).

#### 4.1.3 Kidney Proximal Convoluted Tubule Epithelial Cell

As a second example cell type, we also show the results for kidney proximal convoluted tubule epithelial cells in 10 mice of 1 month, 3 months, 18 months and 30 months. The model has an R-squared value of 0.8 during training and testing (Figure 4a). These cells line the proximal convoluted tubules and enable reabsorption of water, nutrition and electrolytes to facilitate transport into the peritubular capillaries.

**Figure 4:**
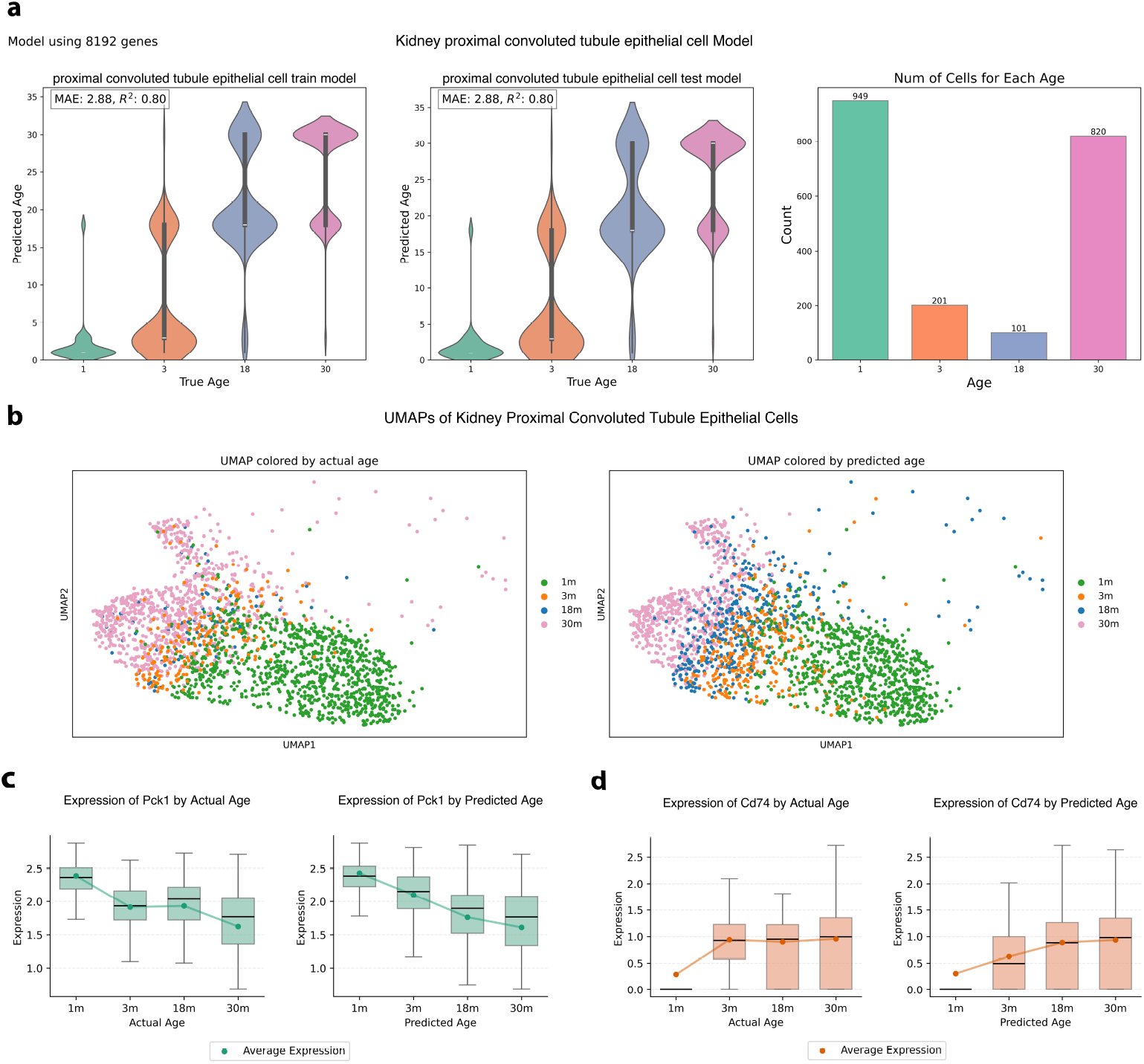
TMS Kidney Proximal Convoluted Tubule Epithelial Model. **a**: Violin plots of true age versus predicted age for kidney proximal convoluted tubule epithelial cell training data (left) and testing data (middle), along with a bar plot showing the number of cells at each age in training and test sets (right). **b**: UMAPs colored by actual ages (left) and predicted age (right). **c**: Box plots of PCK1 expression across actual ages (left) and predicted ages (right), with dotted trend lines indicating average expression change with ages. **d**: Box plots of CD74 expression across actual ages (left) and predicted ages (right), with dotted trend lines indicating average expression with age.

The UMAP visualization of the cells colored by actual ages displayed a largely continuous trajectory from 1-month to 24-month cells, although 18 months were mixed with those of other ages (Figure 3a). By contrast, the UMAP colored by predicted ages revealed a better age separated trajectory (Figure 3b). Notably, a subset of 30-month cells was predicted to be 18 months, suggesting their transcriptional features might be more similar to those of younger cells. Conversely, some cells from the 3-month group were predicted to be 18-months old, indicating that despite sharing the same chronological age, their transcriptomic profiles were shifted toward an older state. These patterns suggest that the predicted ages capture variability that is not visible when cells are grouped only by the donor age.

To investigate whether our model identifies genes associated with the reabsorption differences observed during aging, we identified among the top 100 ranked genes by correlation with predicted age those associated with this cell type. One of our focuses is *PCK1* ranking at 8th with correlation value of −0.50. *PCK1* (phosphoenolpyruvate carboxykinase) catalyzes the conversion of oxaloacetate to phosphoenolpyruvate, thereby maintaining blood glucose homeostasis[21]. Both overall and average expression levels exhibited a clear declining trend with actual age, and a steadier and more consistent decline when stratified by predicted ages (Figure 4c).

Another gene we focused on was CD74, a chaperone gene for the major histocompatibility complex (MHC) class II molecules, that is responsible for presenting foreign peptides to CD4+ T cells. In the proximal convoluted tubule model, CD74 ranked 35th among 8,192 genes with a Pearson correlation of 0.40. Previous work has shown that *CD74* is associated with kidney tubule inflammation[22]. In our TMS dataset, CD74 expression displayed a stepwise pattern across chronological ages. Box plots and average expression values demonstrated an overall upward trend with actual age (Figure 4d). When stratified by predicted ages, the increase appeared more monotonic (Figure 4d), indicating that our model leads to the identification of genes that change monotonically with predicted age.

#### 4.1.4 Conserved age associated genes across tissues

To identify genes that exhibit consistent age-associated transcriptional changes across cell types, we aggregated the Pearson correlation results of the predicted ages and applied the Benjamini–Hochberg correction to obtain adjusted *p*-values (see Methods). Using a significance threshold of *p*_*adj*_ *<* 0.05, we identified 17,090 genes combined across cell types (S Figure 10). Genes were ranked by their cumulative significance, sum of log-adjusted *p*-values, across all cell types and organs, and we focused on the top 100 genes (Figure 5a). The majority (68 genes) were ribosomal genes. In addition, there are 8 transcription or stress related genes, *DBP, H3F3B, TPT1, MALAT1, PTMA, DDX5, RBM3, EEF2*. Several genes linked to inflammation and immune response showed strong age associations across tissues, including the S100 family, *S100A6, S100A8, S100A9* and *S100A11*, and other interferon-related genes *B2M* and *IFITM3*, highlighting the broad involvement of innate and adaptive immune pathways in age-associated transcriptional shifts[23]. Another broad category of genes our analysis captured are associated with the cytoskeleton, which is related to cell movement and structure: *PPIA, PFN1, TMSB4X, TMSB10, ACTB, CFL1*. We also observed contributions from extracellular matrix /adhesion, iron metabolism, histocompatibility complex proteins and housekeeping… (Figure 5b&supplementary table 1).

**Figure 5:**
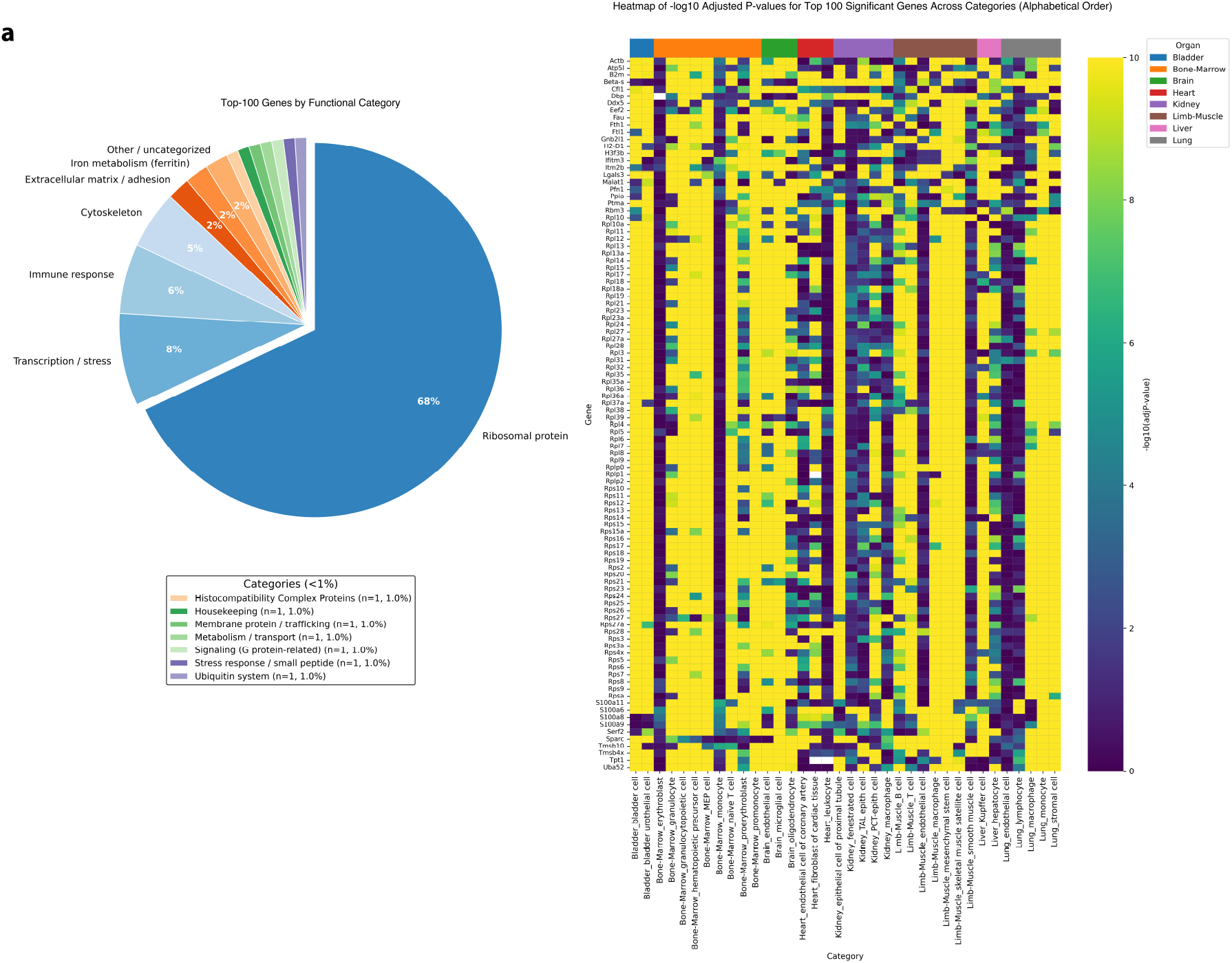
Top 100 Most Significant Gene Analysis. **a**: Pie chart summarizing the functional categories of the top 100 age associated genes. **b**: Heatmap showing the top 100 genes in alphabetical order highlighting in a gradient color scale with lighter color (yellow) depicting the higher log significance value and darker color (blue) depicting the lower log significance value.

We also identified genes that are consistently age associated across cell types using a second approach involving differential gene expression (DEG) analyses in predicted age groups across all cell types and organs. Again, we used *p*_*adj*_ *<* 0.05 as the cutoff point to identify genes with robust age-associated expression patterns. Across all comparisons, 1288 genes were identified in the predicted age DEGs. We focused on the top 20 most significant genes for each age group, which resulted in 93 genes in total (Figure 6a). We observed some shared and condition-specific markers, offering insight into which genes are consistently captured by our predictive model. This unique sets of predicted-only genes potentially represent subtle molecular features that our model detects but may be overlooked in direct age-based comparisons.

**Figure 6:**
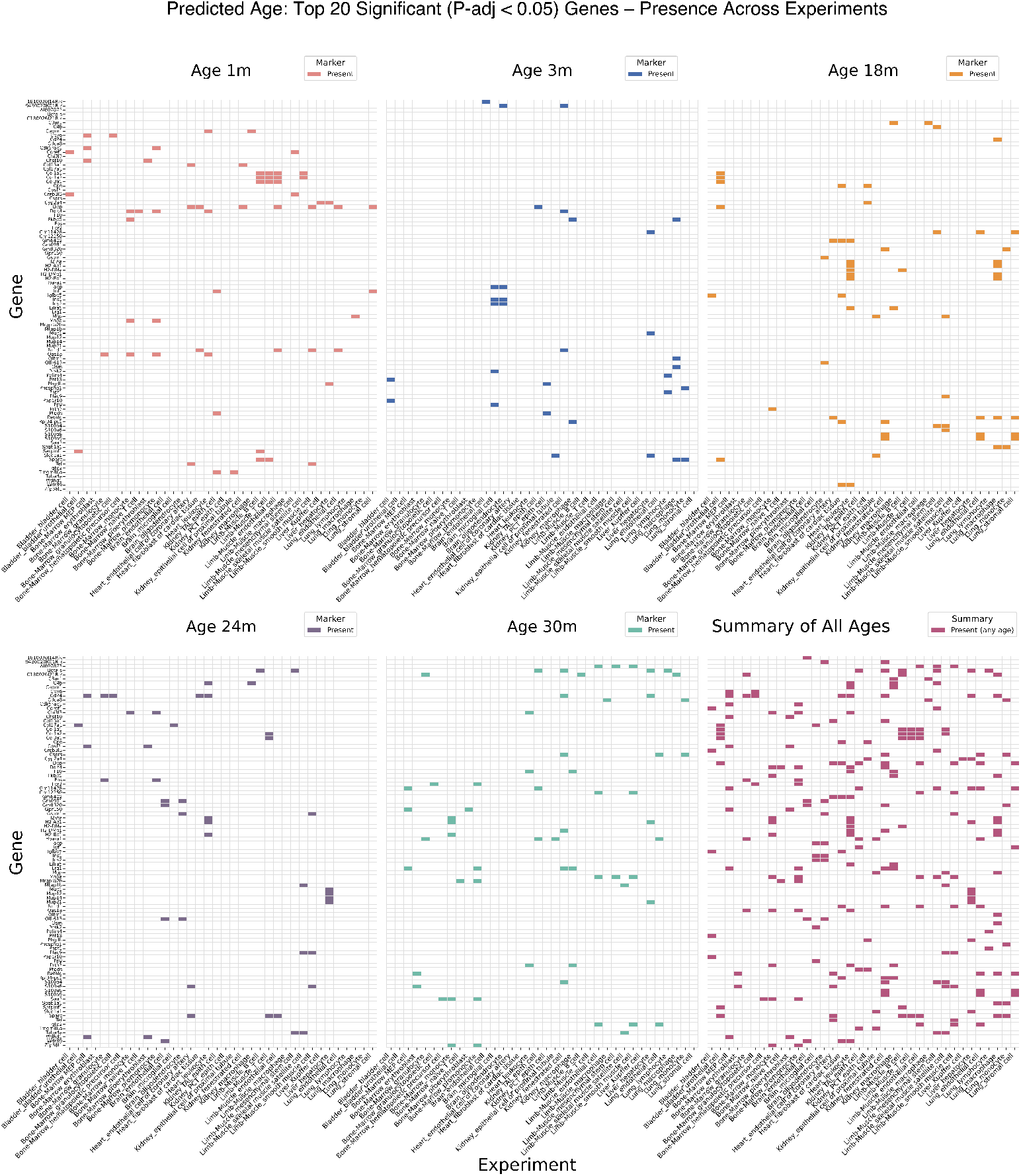
Top Differentially Expressed Gene Analysis. **H**eatmaps showing the top 20 significant genes of each age group for DEG analysis across predicted ages. Color shades represent the presence of the significance value from the DEG analysis in specific cell type. Different colors represent different age groups.

By examining the functional composition of the top genes from the DEG analysis, we again recognized a significant proportion of immune response genes (Figure 7a), consistent with the correlation based analysis. Several immune-related genes were identified, including members of the S100 family (*S100A6, S100A8, and S100A9*) (Figure 7b&c). The DEG analysis did not capture *S100A11*, but it captured a major histocompatibility complex class II gene *CD74*, the gene we mentioned previously while presenting our kidney proximal convoluted tubule epithelial cell model, and some other genes that play important roles in varied immune process. We also observed the presence of transcription and stress genes (see supplementary table 2). Extracellular matrix /adhesion genes were also identified in this analysis (see supplementary table 2) along with a few histocompatibility genes, *H2-Aa, H2-Ab1, H2-DMa, H2-DMb1 and H2-Eb1*. In total, there are 6 genes overlapped between the correlation and DEG analyses (Figure 7c).

**Figure 7:**
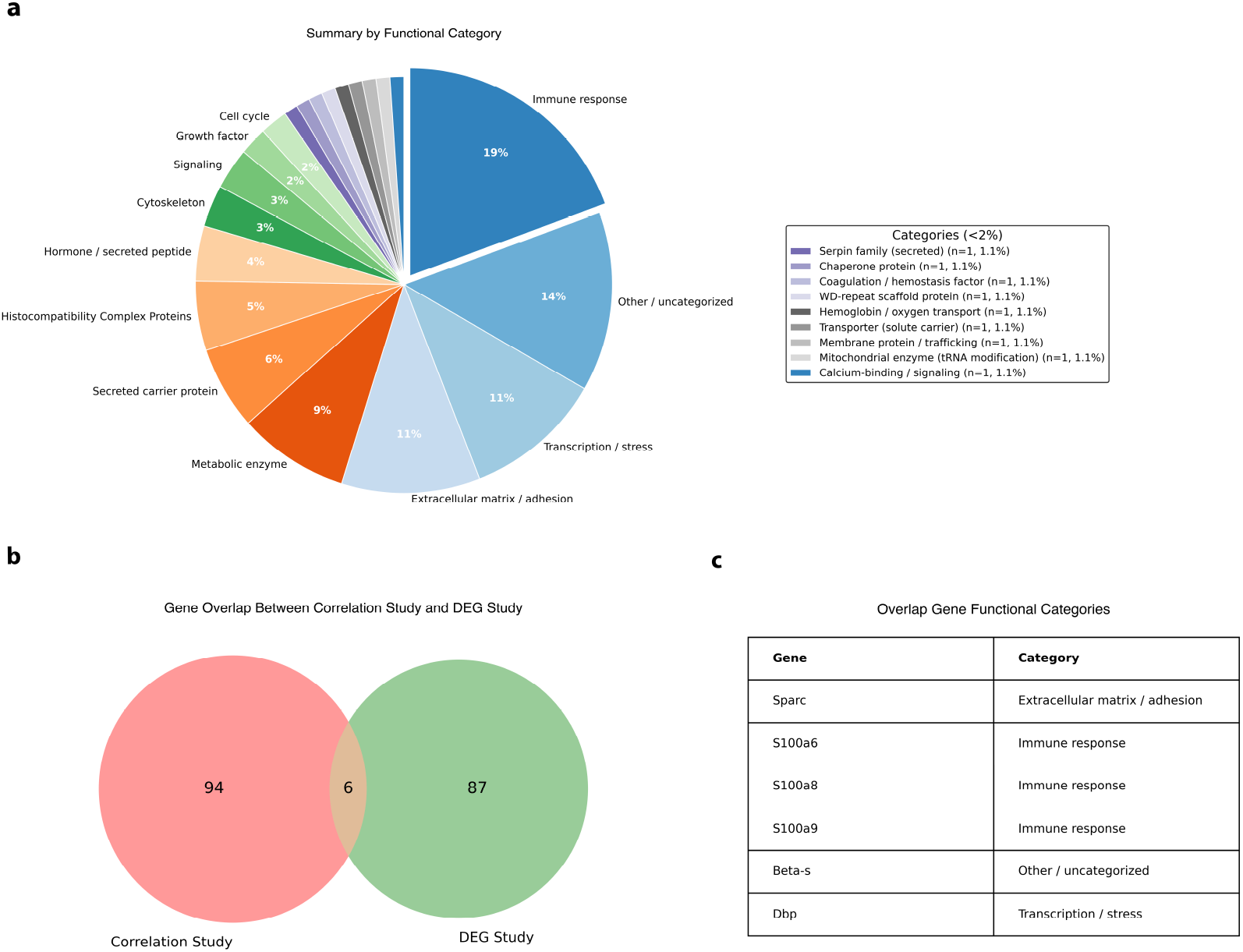
Top Differentially Expressed Gene Classifications and Overlap Analysis. **a**: Pie chart summarizing the functional classifications of the same genes in Figure 6. **b**: Venn diagram showing the overlap of significant genes between the correlation and DEG analyses. **c**: Table showing the overlapping genes

## 5 Discussion

In this study, we present a new scRNA aging clock, *scMLEAge*, based on scRNA transcriptomic gene counts. Our framework builds cell-type–specific regression models of gene expression frequencies and then estimates the probability of a cell to be associated with each age by considering the probability of the counts given the expected gene expression frequencies. The final age prediction for a single cell is obtained by evaluating the likelihood of all possible age groups and selecting the age with maximum probability. *scMLEAge* incorporates a hyperparameter to flexibly select an optimal number of genes to include in the analysis. This approach serves as a useful tool for explicitly modeling the count-based nature of scRNA-seq data.

The implicit assumption in our method is that the cells of an individual have a range of cellular ages that are not completely determined by the age of the individual from which they originate. Thus cellular ages are heterogeneous within an individual, and the cells within a single individual can be partitioned based on their estimated cellular age. The caveat is that we do not know the age of individual cells a priori and so use the age of an individual to train our models, assuming that most of the cells will be approximated by the chronological age of the individual. Despite this training approach, the model generates a range of cellular ages for a single individual, and these can then be used to identify markers of cellular ages.

Compared with regression-based approaches such as ElasticNet, *scMLEAge* provides several important advantages. ElasticNet assumes linear relationships between normalized gene expression and chronological age, which can be effective in bulk or well-averaged data but is less suited to the sparsity and overdispersion inherent to scRNA-seq. Rather than fitting into a regression trend, *scMLEAge* models the raw count data directly within a Poisson and maximum likelihood framework, thereby aligning more closely with the stochastic nature of transcriptional events at the single-cell level. This generative approach allows us to incorporate sequencing depth explicitly, produce interpretable age-group frequency estimates, and capture cell type–specific heterogeneity. Benchmarking on limb muscle cell types from the TMS dataset, *scMLEAge* consistently achieved higher *R*^2^ values than ElasticNet in most cell types, demonstrating that *scMLEAge* not only improves predictive accuracy but also offers a principled framework that reflects the biological properties of single cells, while also being substantially more computationally efficient than ElasticNet.

Although scRNA-seq data are often described as overdispersed relative to a Poisson distribution, several aspects of our framework mitigate this concern. First, our model operates on gene expression frequencies aggregated within age groups and cell types, which reduces cell-level variability and stabilizes the mean–variance relationship. Second, the likelihood is evaluated jointly across a large number of genes, such that deviations from the Poisson assumption at individual genes have a limited impact on the overall age estimate. In this setting, the Poisson model serves as a computationally efficient and well-calibrated approximation for capturing the dominant signal in count-based data. Consistent with this, we observe strong empirical performance across multiple cell types, suggesting that strict modeling of overdispersion is not required to achieve accurate age prediction in this framework. Alternative models such as the negative binomial could explicitly account for overdispersion and may further improve performance in certain settings but estimating the dispersion parameter in a cell-type– and gene-specific manner would substantially increase model complexity and was beyond the scope of this study.

While *scMLEAge* performed well across most cell types, a few models exhibited lower predictive accuracy. For example, the clock for bone marrow granulocytopoietic cells achieved an *R*^2^ of 0.17, whereas the model for mature granulocytes had a higher *R*^2^ of 0.67. Granulocytes represent a terminally differentiated population derived from granulocytopoietic progenitors. Previous studies have reported that aging affects mature granulocytes more strongly than their progenitors [24], which may explain the limited age-associated signal captured in our granulocytopoietic cell model.

Muscle stem cells (muscle satellite cells) are well known to decline during aging. In both humans and mice, their abundance decreases with age, which reduces regenerative potential, delays recovery from injury, and contributes to deterioration in posture and mobility [25, 26]. Our model captures transcriptional changes in limb-muscle satellite (stem) cells, achieving an *R*^2^ of 0.8. Consistent with this, the scAge single-cell DNA-methylation clock applied to mouse muscle stem cells (muscle satellite cells) also detected a significant rise in epigenetic age from 1.5 to 26 months[27], reflecting measurable aging in muscle stem cells. To further explore the genes contributing to this model, we examined features ranked highly by our model. One example is *COL6A1*, a collagen VI gene essential for extracellular matrix integrity. Previous studies found that aging causes the loss of collagen VI genes, which leads to reduced self-renewal capability and impaired muscle regeneration[28].

Aging of kidney proximal tubules is frequently characterized by inflammation [22]. Our model captured transcriptional changes in kidney proximal tubule epithelial cells, with an *R*^2^ of 0.8. Among the features contributing to this model, we identified *PCK1*, a metabolic gene critical for gluconeogenesis and blood glucose maintenance[21]. It has been reported as a key down-regulated marker in aged kidney cells[29]. Aged kidney tubules also undergo metabolic shifts, DNA damage, and cellular senescence that impair reabsorption, and can further progress to chronic kidney disease. Several of these alterations are related to the major histocompatibility complex class II (MHC-II) genes. In our model, one such MHC-II gene was *CD74*, an immune gene that associated with kidney tubule inflammation. Expression patterns of both *PCK1* and *CD74* in the TMS dataset aligned with previously reported age-related changes, supporting that our model generates biologically meaningful signals to predict cell-level aging trajectories.

On a broader scale, we also examined expression changes conserved across multiple cell types and organs in predicted ages. Among the top age-correlated and differentially expressed genes, the majority were ribosomal genes and immune response genes. Although high expression of ribosomal genes is sometimes interpreted as an indicator of low-quality or stressed cells, in our analysis we retained these features, as they may instead reflect biologically relevant stress responses or transcriptional signatures of dying cells. These genes encode structural proteins that form the ribosome, and their abundance has been reported to decline with increased DNA methylation during aging [30, 31]. The presence of ribosomal genes among our top features underscores the central role of protein synthesis and translational regulation in the aging process. To assess whether our findings were driven primarily by ribosomal genes, we additionally rebuilt and evaluated the models after removing ribosomal genes from the feature set. All model performance were unchanged (S Figure 11), indicating that the age-predictive signal is not dependent on ribosomal genes. This comparison supports that the other age-associated markers identified in our analysis capture genuine aging-related transcriptional programs.

Both of the methods we used to identify age associated genes across multiple cell types found immune-related S100 family genes (*S100A6, S100A8, and S100A9*) and many other antimicrobial factors, consistent with prior studies showing that aging is accompanied by persistent activation of immune responses [32]. The study also identified *SPARC*, a matricellular glycoprotein secreted into the extracellular niche that regulates cell adhesion, angiogenesis, growth factor binding, and differentiation[33], whose expression has been broadly associated with age-related tissue remodeling and functional decline in mammals. These observations are broadly consistent with prior analyses of the Tabula Muris Senis dataset [8]. In particular, our identification of conserved immune- and stress-associated markers across diverse cell types agrees with this report of common aging-related pathways with 28 overlapping genes (S Figure 12), whereas the variation in predictive performance and top-ranked features across individual cell types is also consistent with prior studies showing that aging signatures can differ, and in some cases even be oppositely regulated, depending on cellular context. Together, these comparisons suggest that *scMLEAge* recovers known aging signals in the TMS resource while also providing a complementary framework for resolving both conserved and cell-type-specific aspects of aging.

Together, these results demonstrate that our framework reliably captures a wide spectrum of age-associated genes, spanning processes such as ribosomal function, immune activation, and other pathways implicated in tissue aging. By integrating signals across diverse biological programs, *scMLEAge* provides a comprehensive and biologically grounded representation of cellular aging. Beyond improving predictive accuracy, this approach offers a scalable and interpretable tool to investigate how aging manifests across tissues, to identify both conserved and cell type–specific mechanisms, and to generate hypotheses about molecular drivers of functional decline. As such, *scMLEAge* has the potential to become a broadly useful resource for dissecting the biology of aging at single-cell resolution.

## 6 Limitations and Future Direction

Although *scMLEAge* provides improved predictive performance compared with regression-based approaches, several limitations remain. First, our current hyperparameter scheme relies on a power-of-two selection of the most age-correlated genes. While this strategy is simple and efficient, it may still include a substantial number of noisy or redundant features, as gene expression values are often correlated and not fully independent. More sophisticated approaches for feature selection or dimensionality reduction—such as stability selection, mutual information metrics, or incorporating prior biological knowledge—could help refine the gene sets and further improve model interpretability.

Second, the current implementation assumes a linear relationship between gene frequencies and age within each group, which does not capture potential non-linear dynamics of aging. Biological aging is rarely a strictly linear process, and incorporating non-linear modeling techniques could allow the model to better reflect age-related transitions and plateau effects across the lifespan.

Third, our framework currently predicts age using discrete age groups defined by the available data. This design limits the model’s ability to interpolate across unobserved intermediate ages. Future extensions could enable estimation of continuous age probabilities or finer-grained age categories, thereby improving temporal resolution and capturing more subtle transcriptional changes that occur between the sampled age groups.

Finally, our study is limited to male mice to avoid potential sex-specific confounding effects. While this simplifies the modeling framework, it may limit generalizability. Extending scMLEAge to include both sexes, and developing sex-stratified or sex-aware models, will be an important direction for future work. Also, extending the model to go beyond the mice datasets to human datasets will be important for evaluating its translational relevance and applicability to human aging.

## 7 Code Availability

The code used to build aging clocks and analysis in the current study is available in the GitHub repository for this paper https://github.com/DaisyCuttie/scMLEAge

## 8 Author Contributions

Conceptualization: C.H. and M.P.; Analysis and figure preparation: C.H. and M.P.; Writing—Original Draft: C.H. and M.P.

