## Supplementary figures and images for "Determining the age of single cells using scMLEAge"

### Supplementary Figure 1

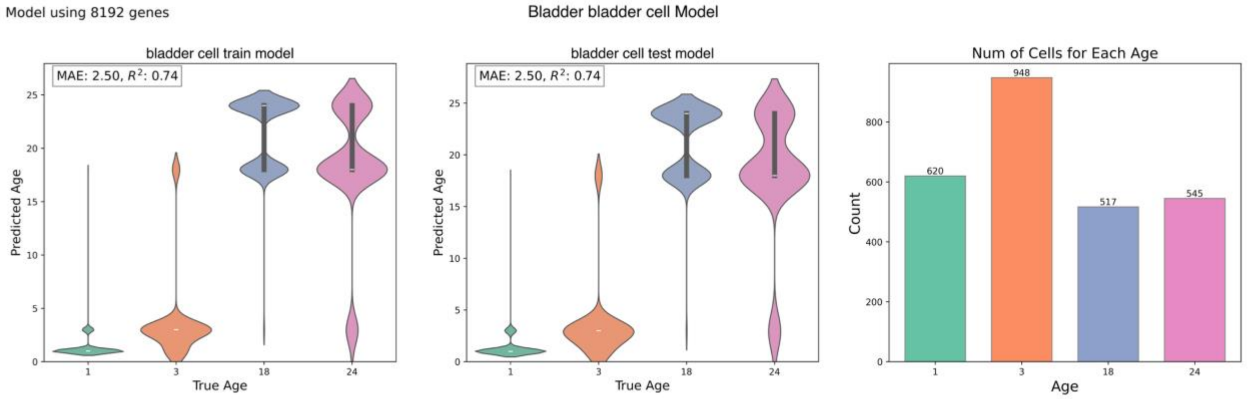

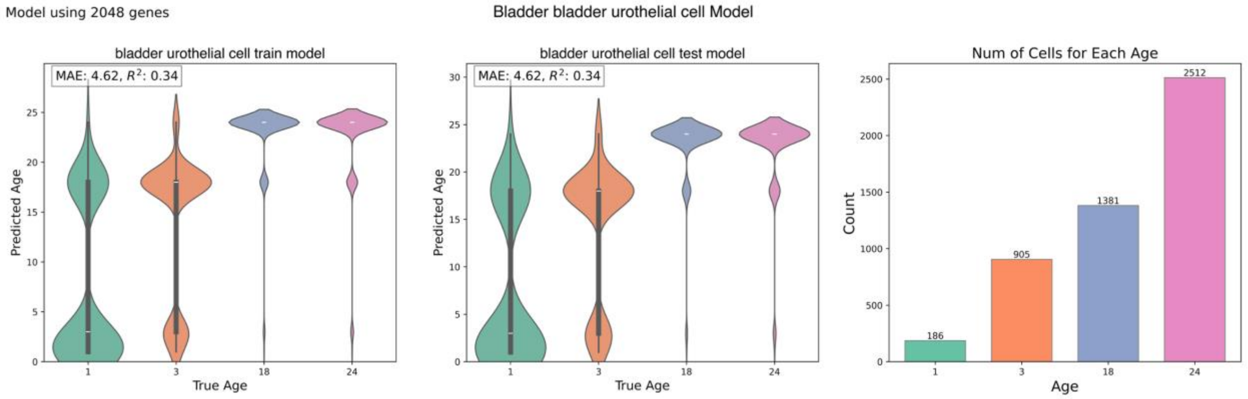

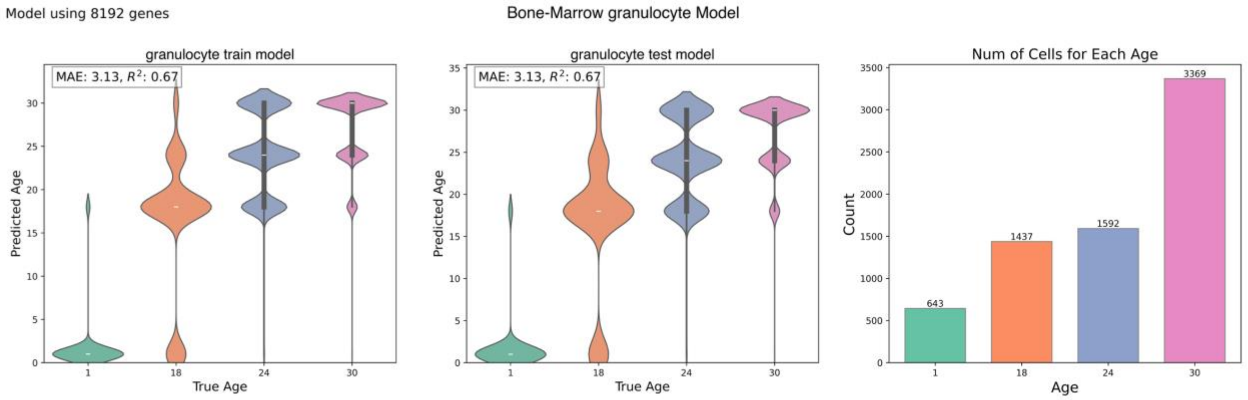

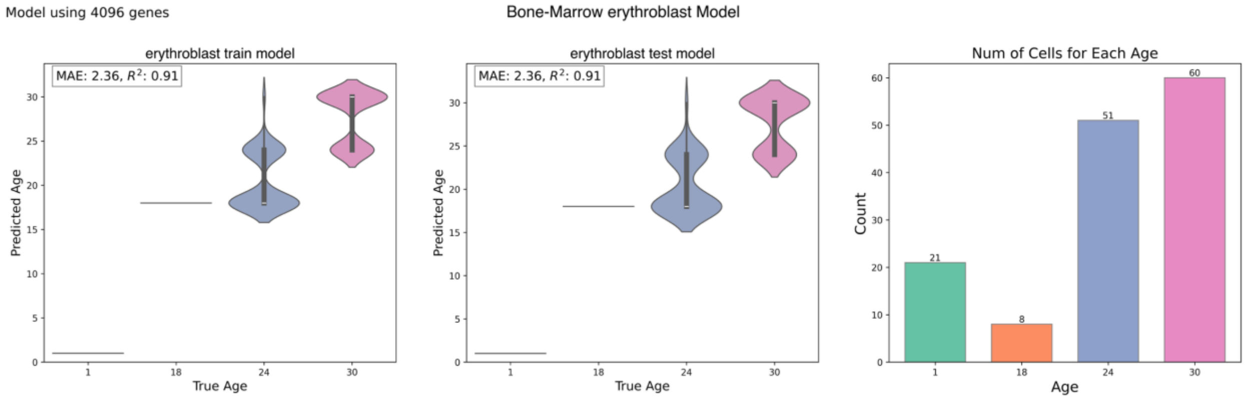

### Supplementary Figure 5

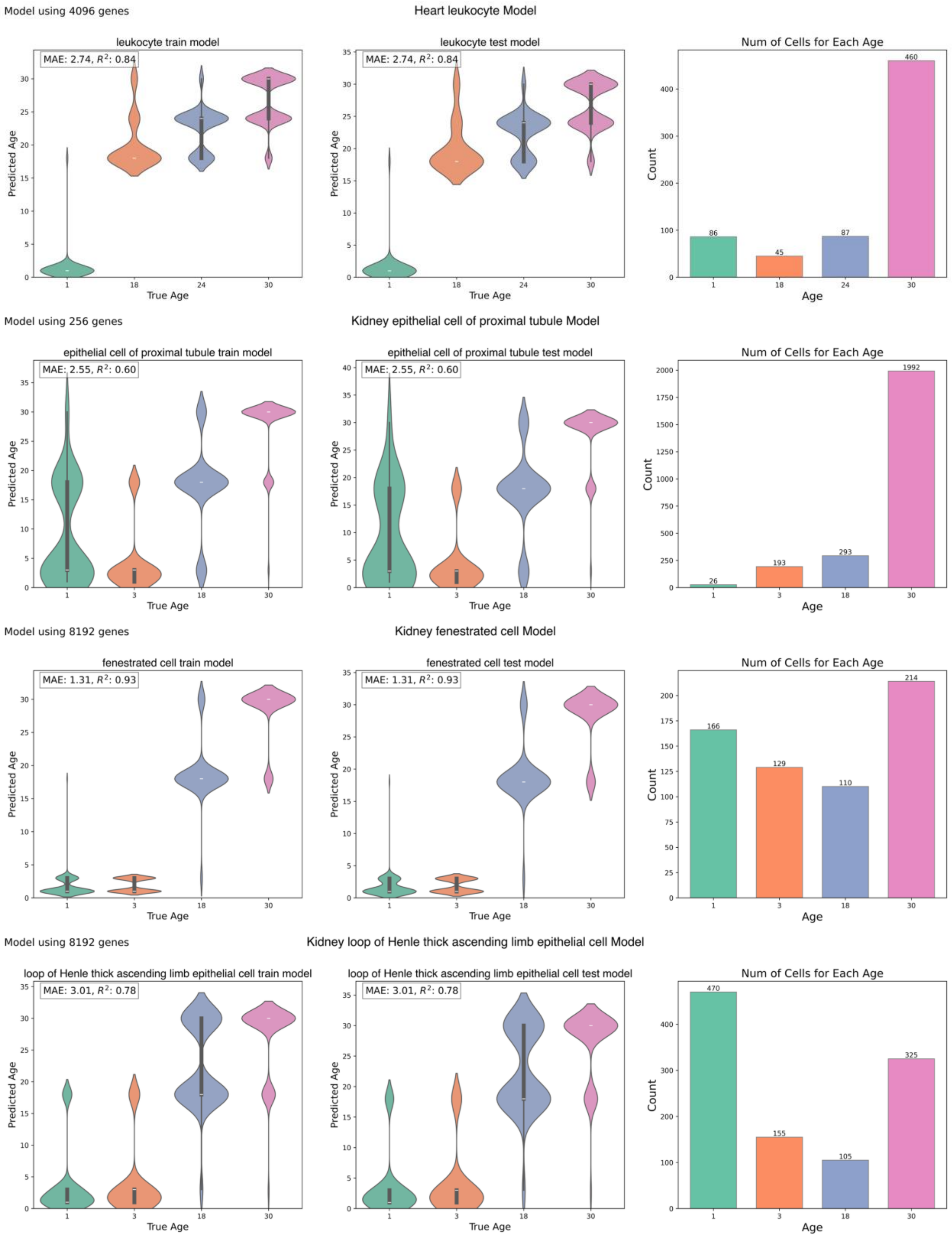

### Supplementary Figure 10

#### Heatmap of Significant Gene Correlation

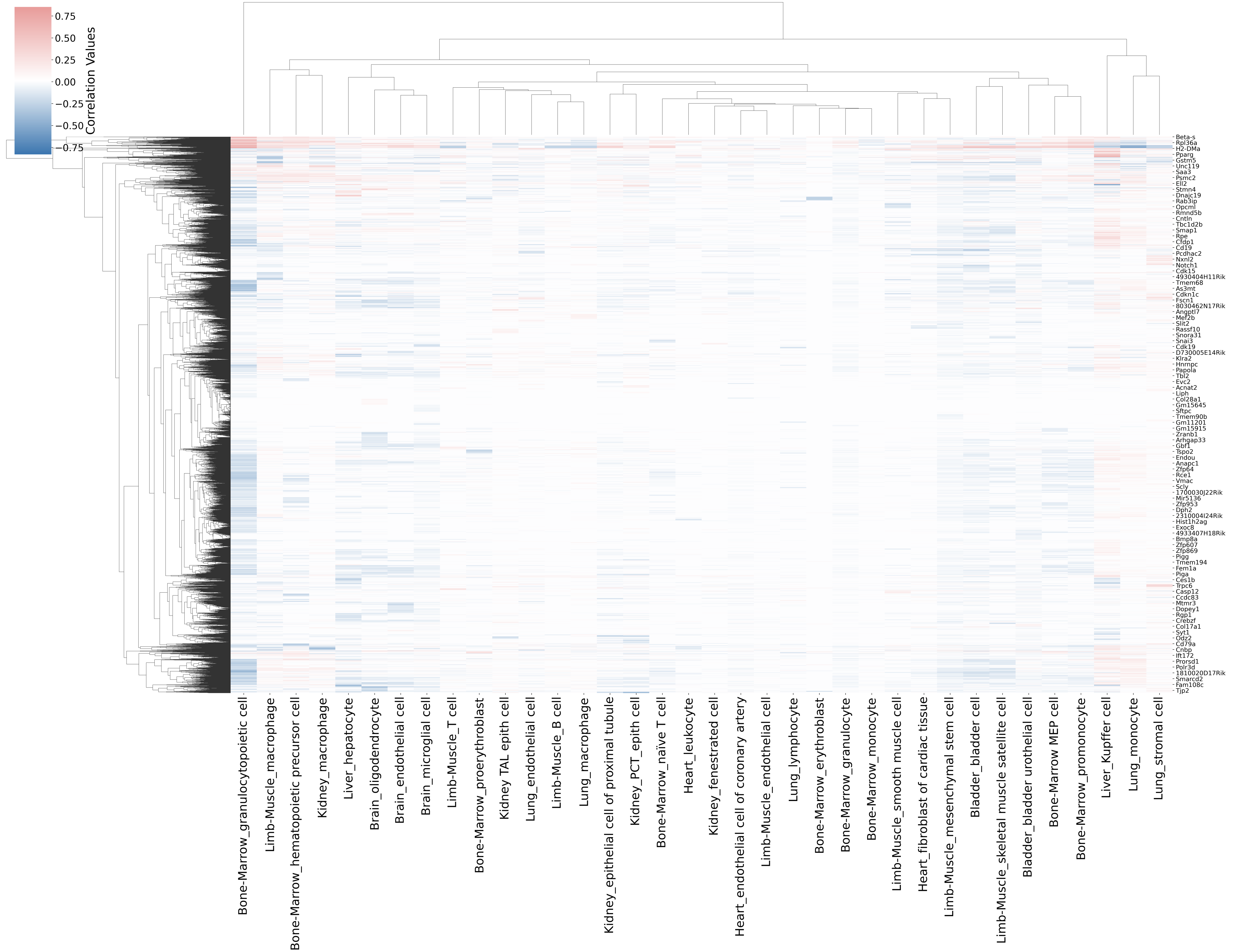

### Supplementary Figure 11

Model Performance Comparison With and Without Ribosomal Genes

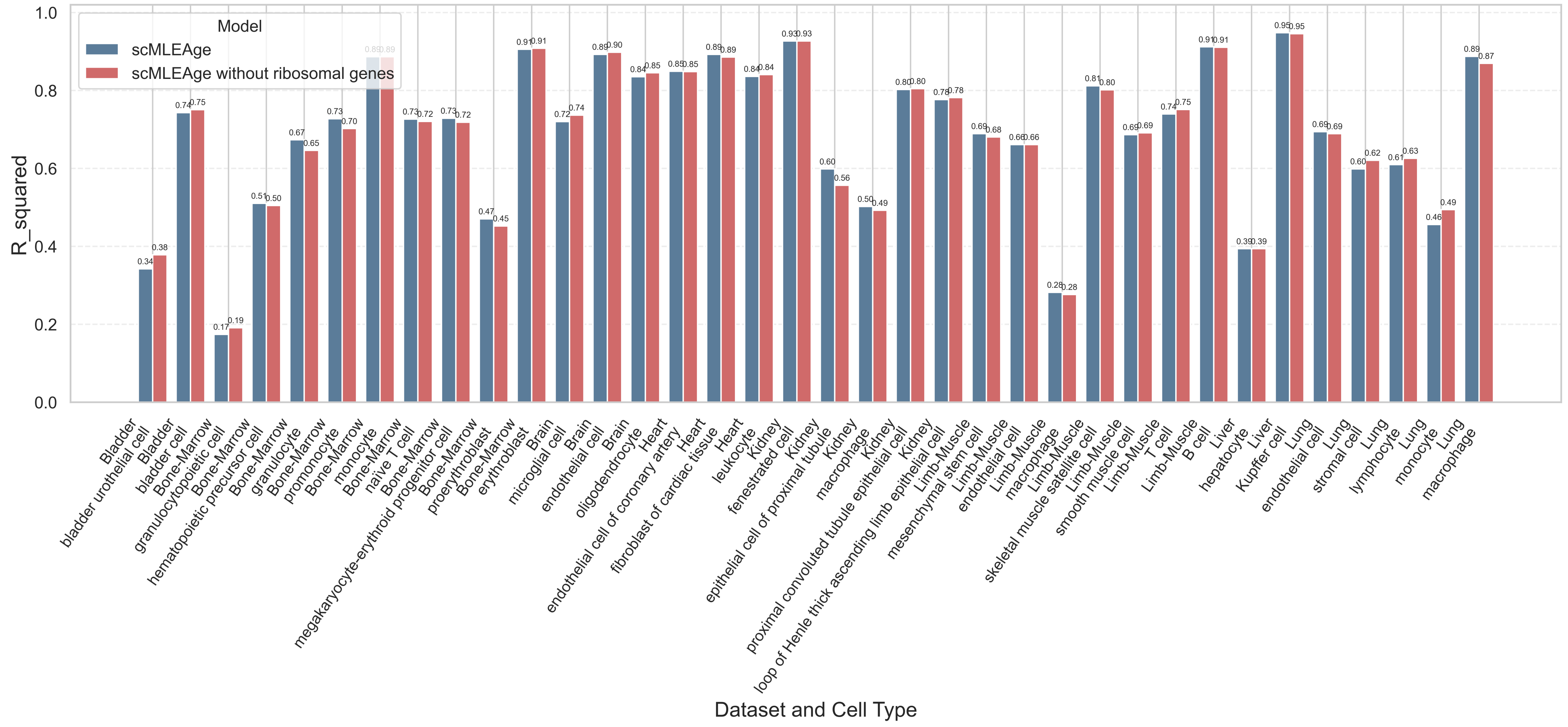

### Supplementary Figure 12

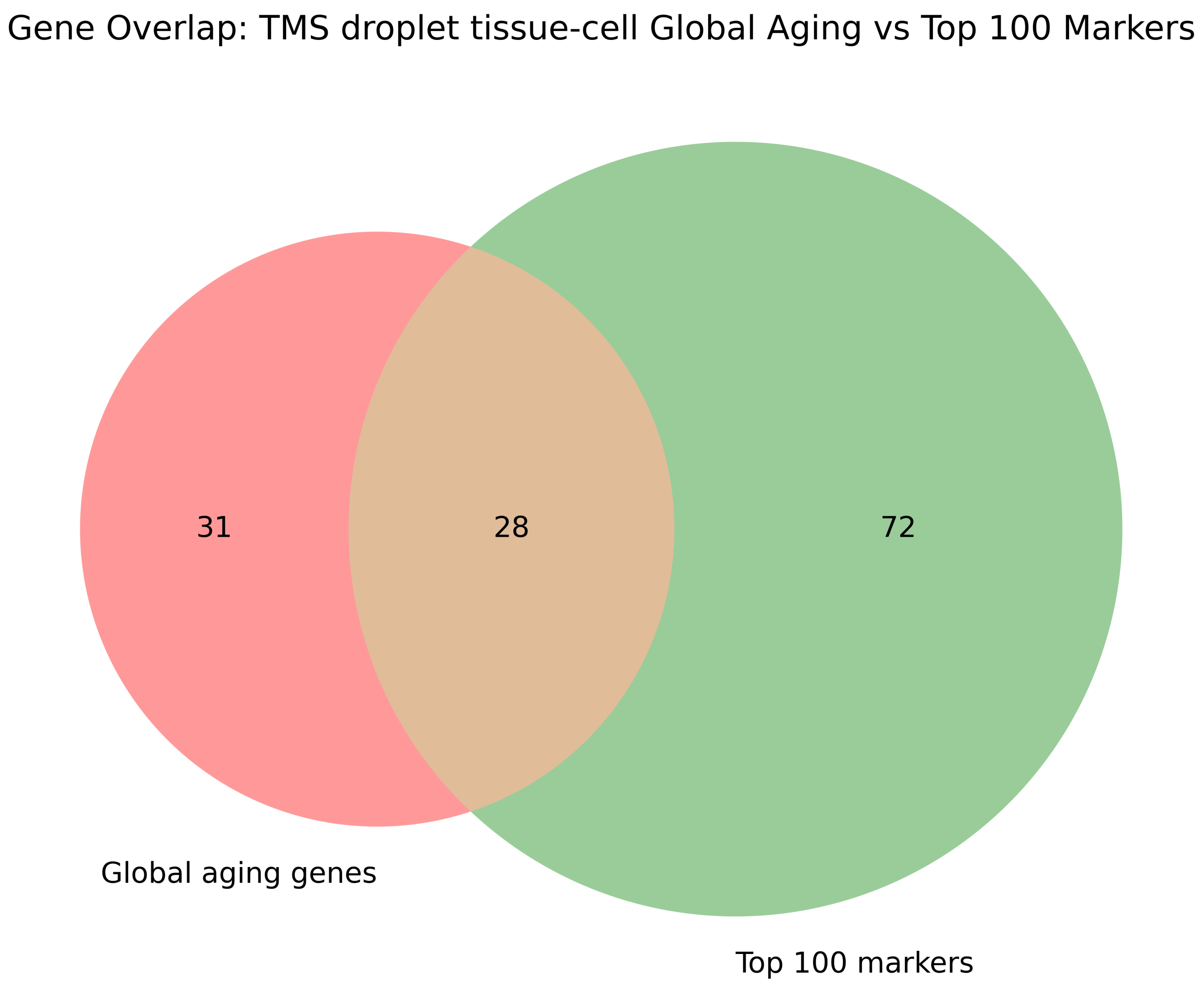
