## Supplementary Figure 2 for "Determining the age of single cells using scMLEAge"

Model using 4096 genes

Bone-Marrow granulocytopoietic cell Model

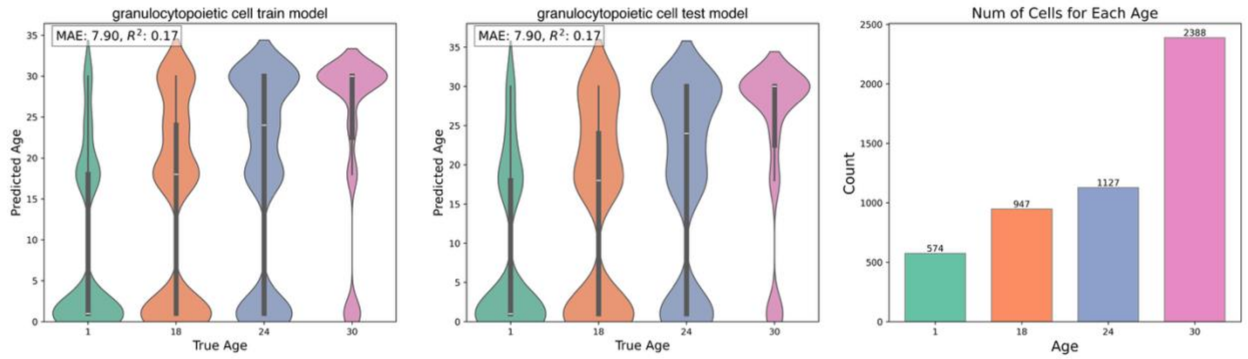


Model using 8192 genes

Bone-Marrow hematopoietic precursor cell Model

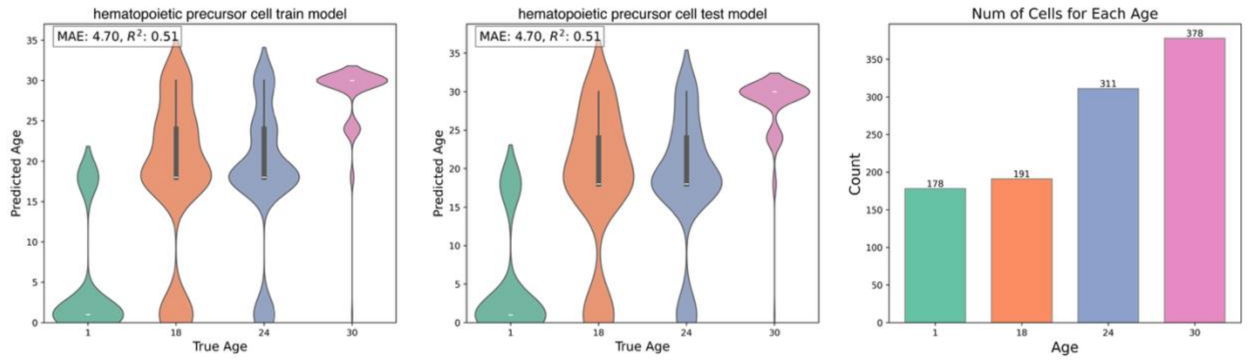


Model using 8192 genes

Bone-Marrow megakaryocyte-erythroid progenitor cell Model

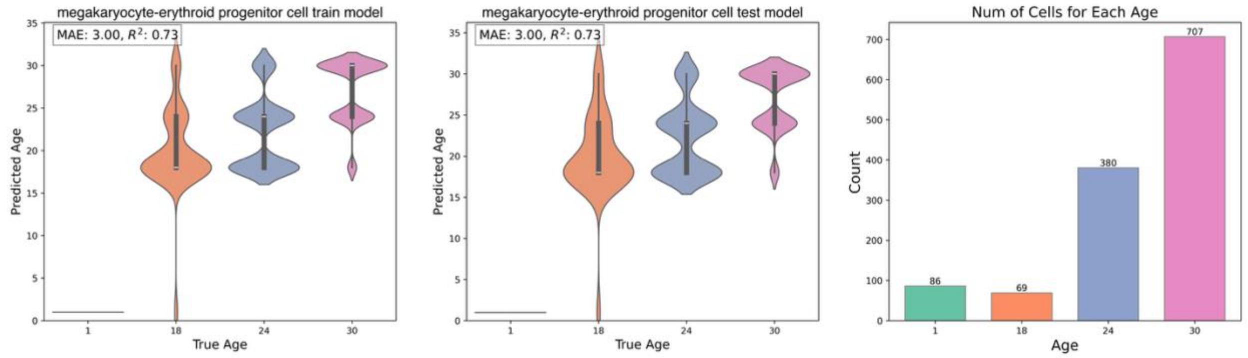


Model using 8192 genes

Bone-Marrow monocyte Model

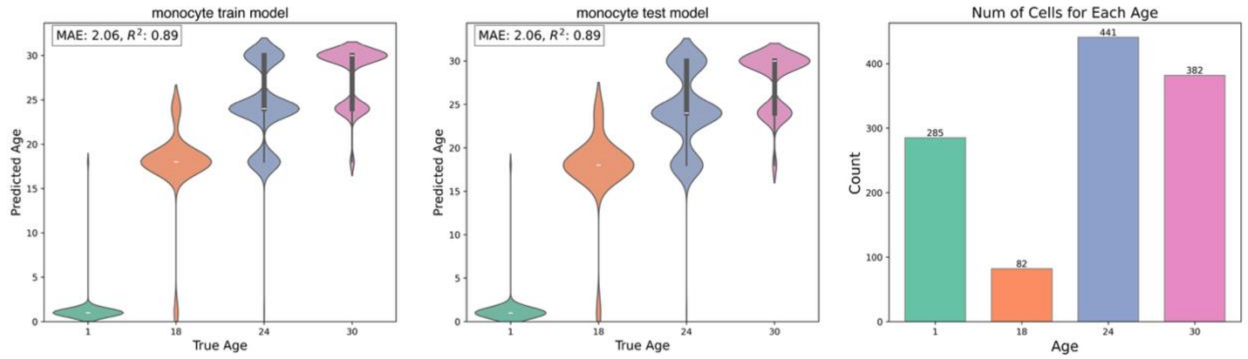
