## Supplementary Figure 3 for "Determining the age of single cells using scMLEAge"

Model using 8192 genes

Bone-Marrow naïve T cell Model

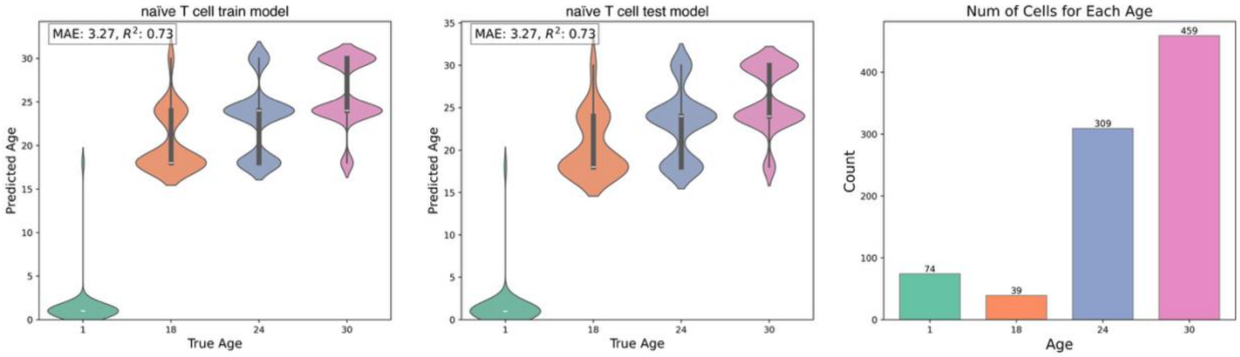


Model using 8192 genes

Bone-Marrow proerythroblast Model

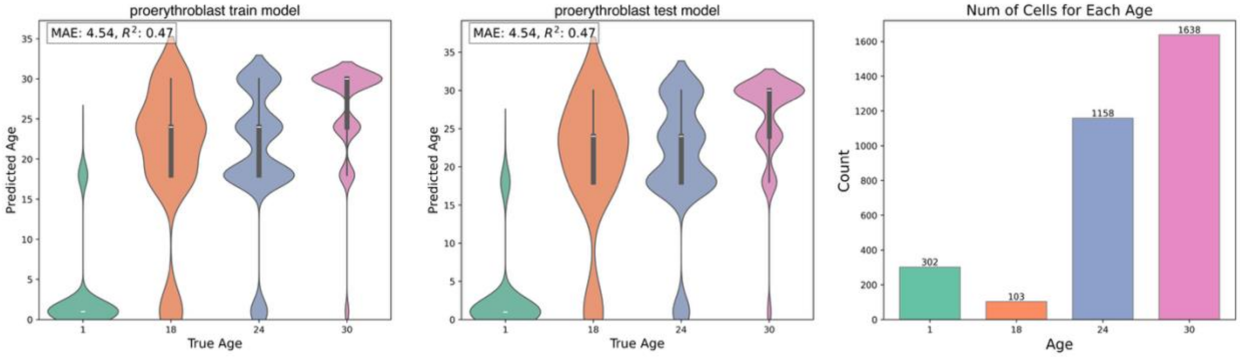


Model using 8192 genes

Bone-Marrow promonocyte Model

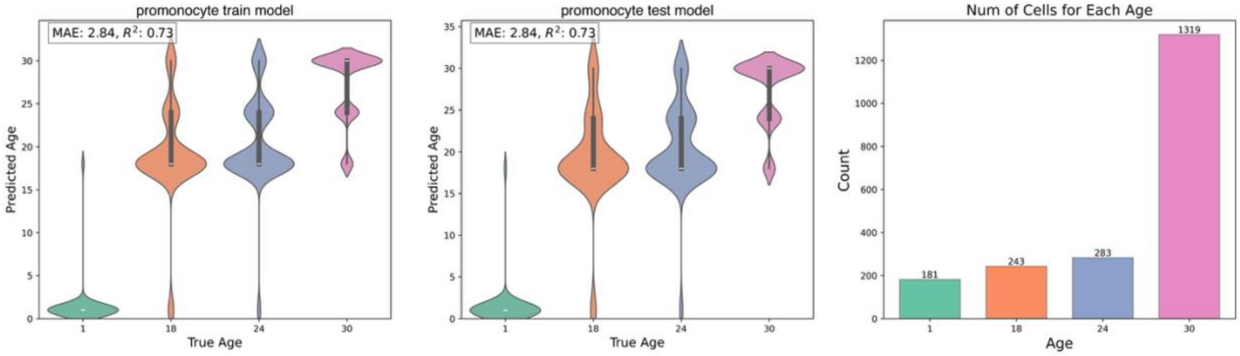


Model using 8192 genes

Brain endothelial cell Model

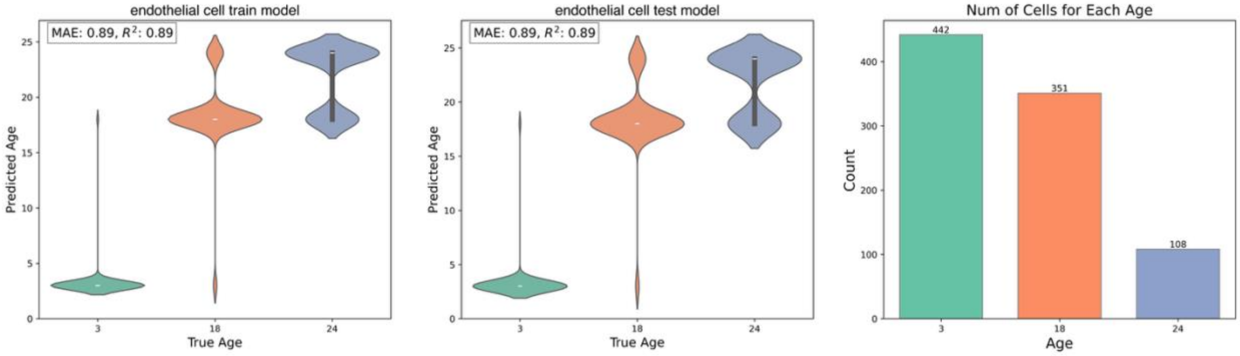
