## Supplementary Figure 4 for "Determining the age of single cells using scMLEAge"

Model using 16384 genes

Brain microglial cell Model

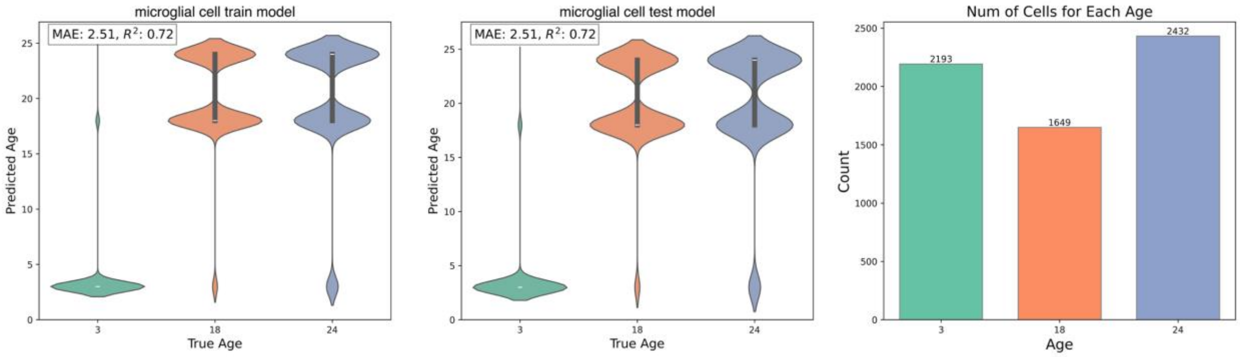


Model using 8192 genes

Brain oligodendrocyte Model

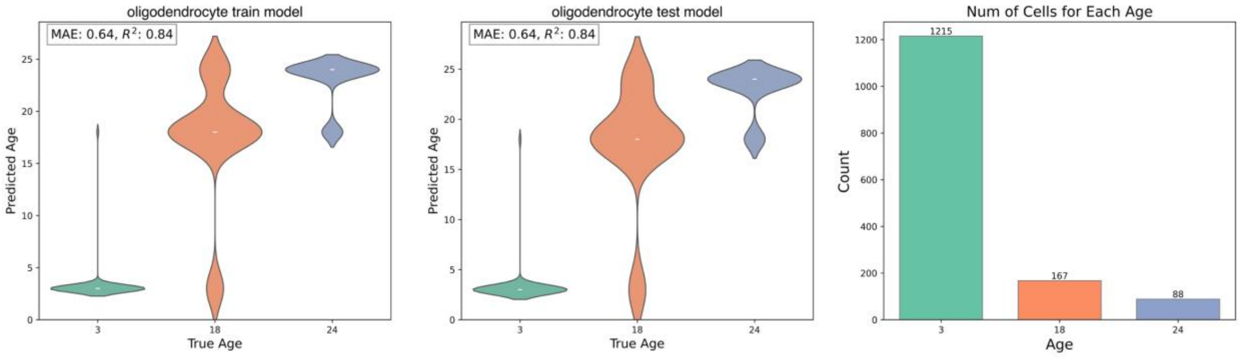


Model using 8192 genes

Heart endothelial cell of coronary artery Model

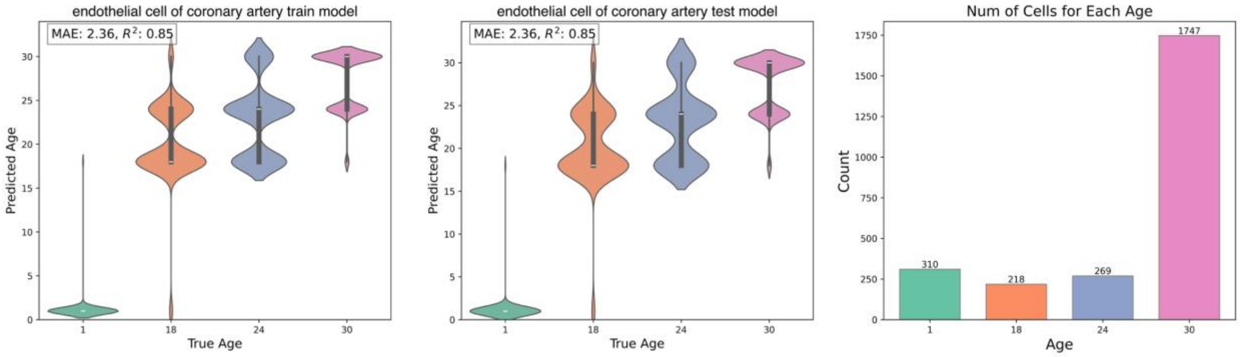


Model using 8192 genes

Heart fibroblast of cardiac tissue Model

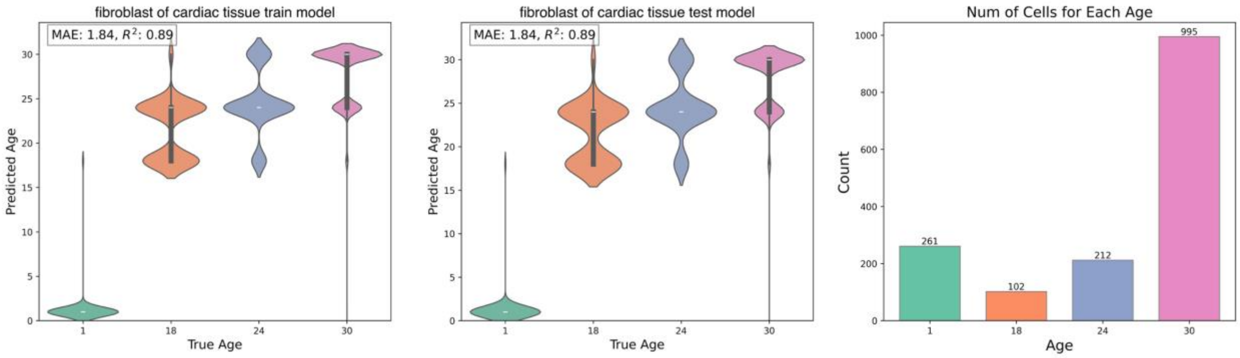
