## Supplementary Figure 6 for "Determining the age of single cells using scMLEAge"

Model using 8192 genes

### Kidney macrophage Model

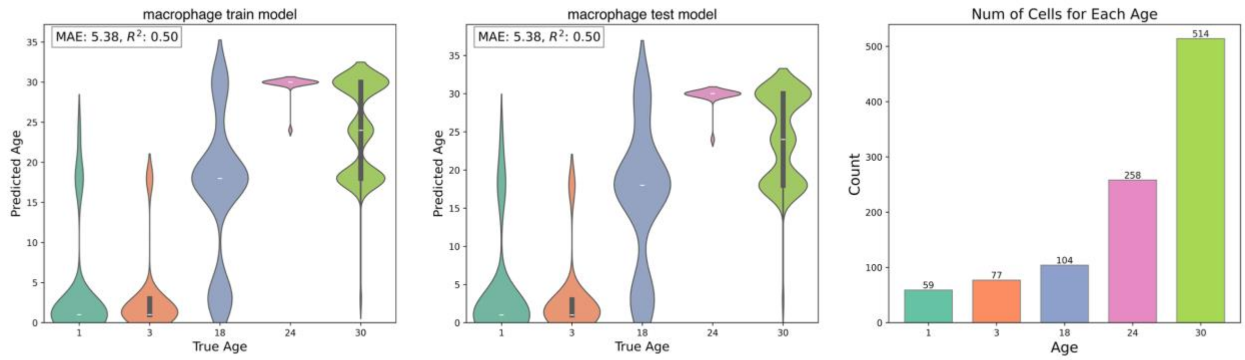


Model using 8192 genes

### Kidney proximal convoluted tubule epithelial cell Model

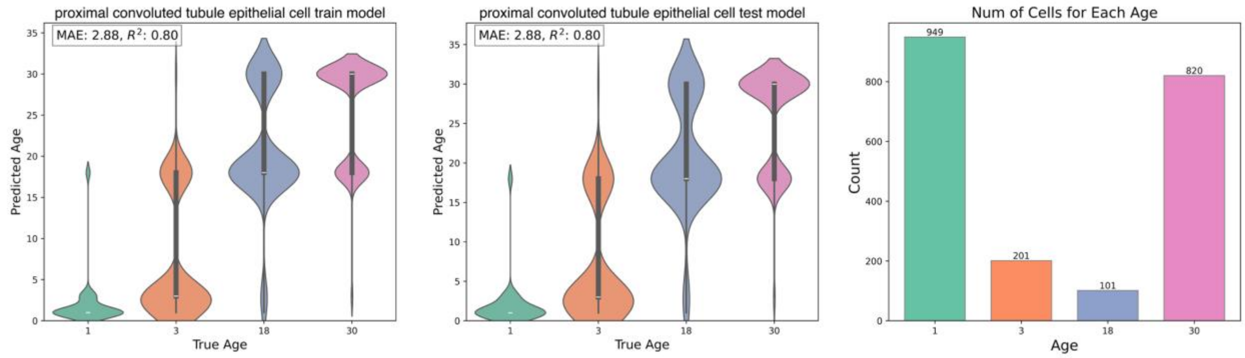


Model using 8192 genes

### Limb-Muscle B cell Model

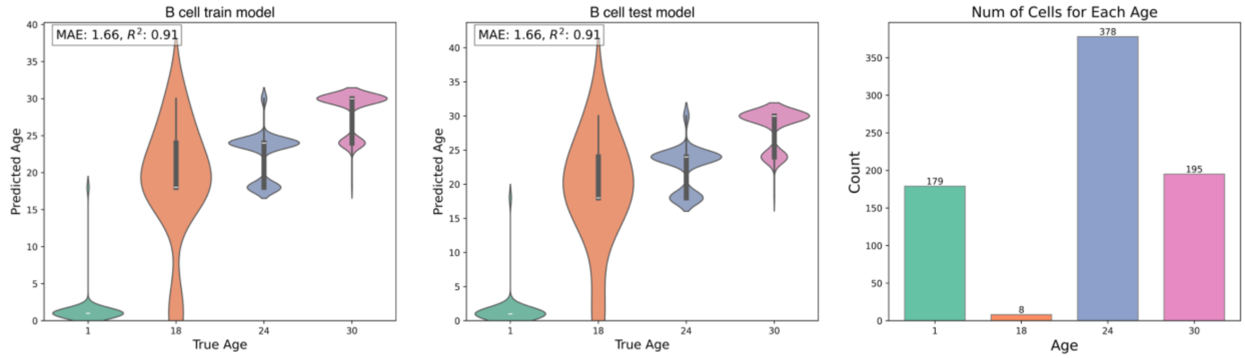


Model using 8 genes

### Limb-Muscle endothelial cell Model

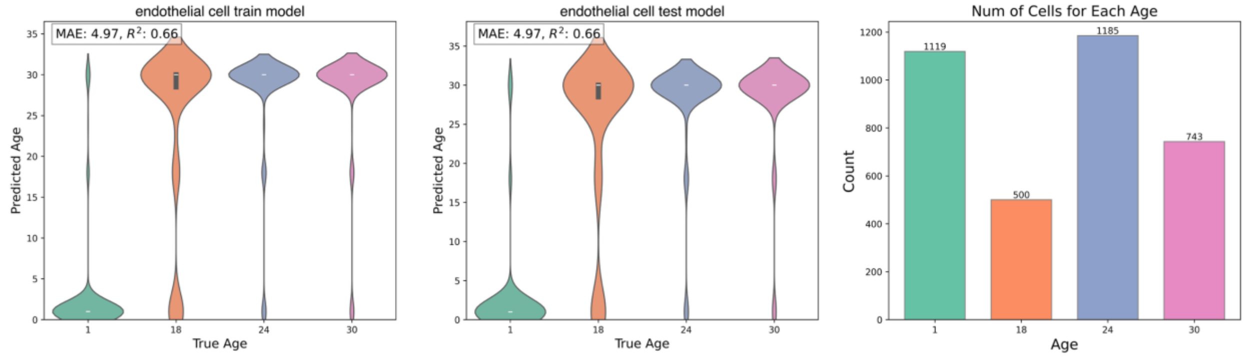
