## Supplementary Figure 7 for "Determining the age of single cells using scMLEAge"

Model using 8192 genes

Limb-Muscle macrophage Model

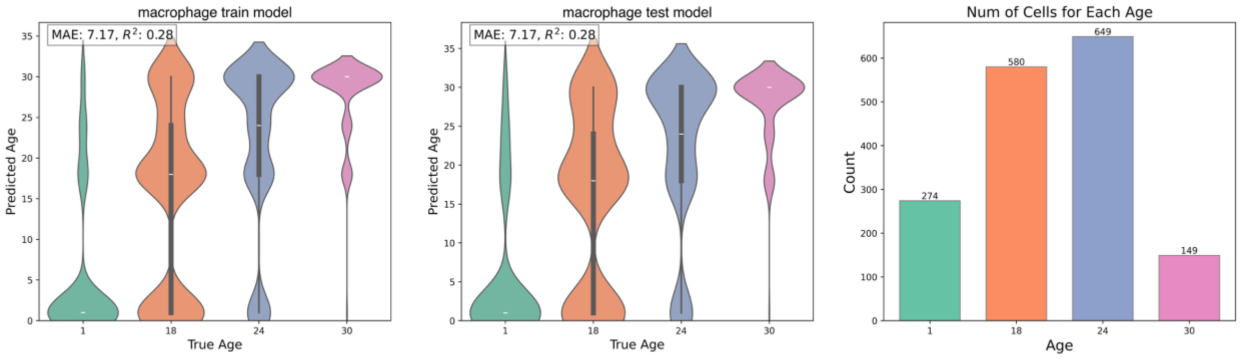


Model using 8192 genes

Limb-Muscle mesenchymal stem cell Model

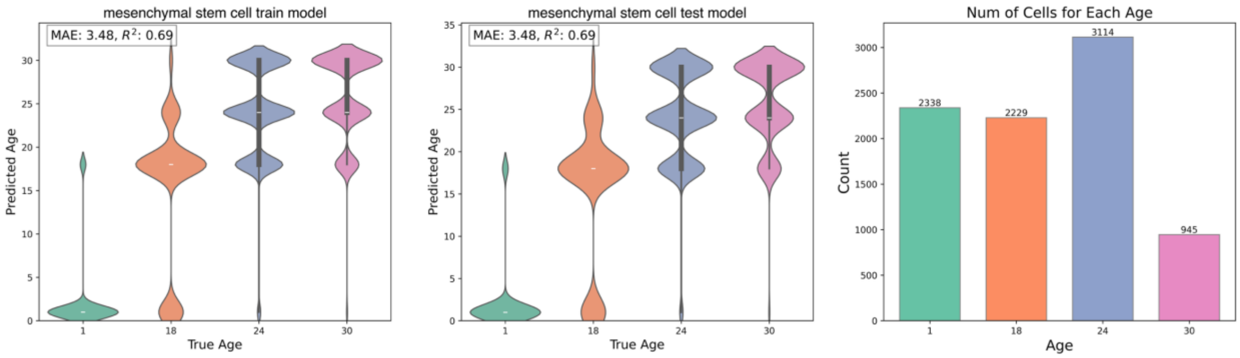


Model using 8192 genes

Limb-Muscle skeletal muscle satellite cell Model

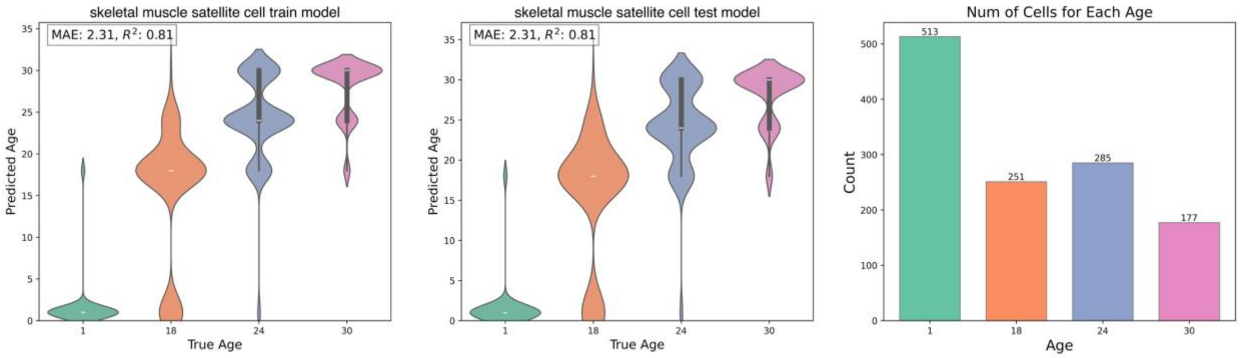


Model using 8192 genes

Limb-Muscle smooth muscle cell Model

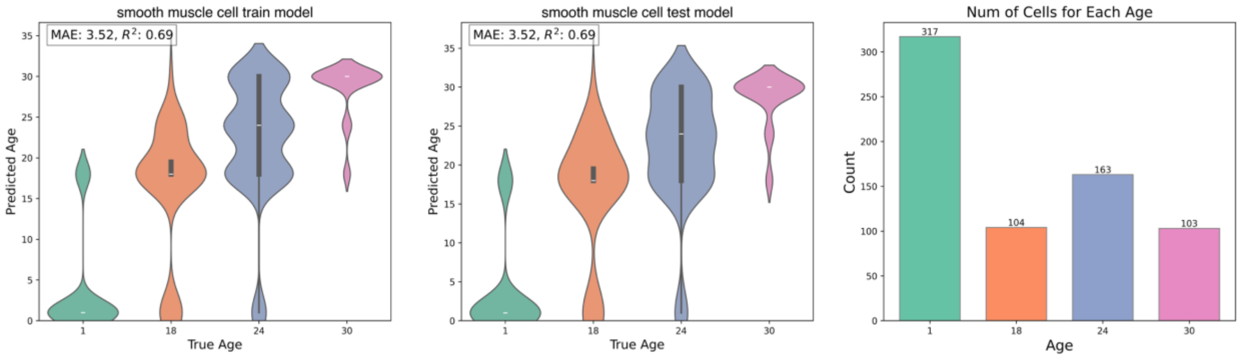
