## Supplementary Figure 8 for "Determining the age of single cells using scMLEAge"

Model using 8192 genes

Limb-Muscle T cell Model

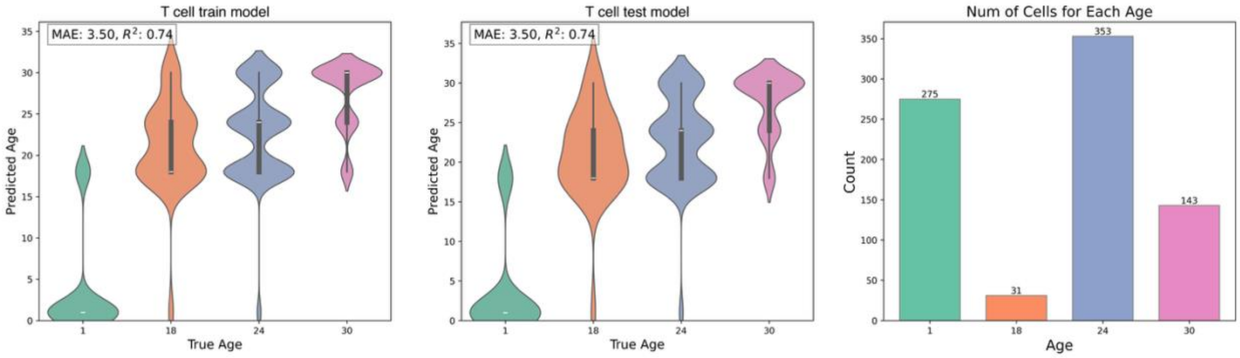


Model using 128 genes

Liver hepatocyte Model

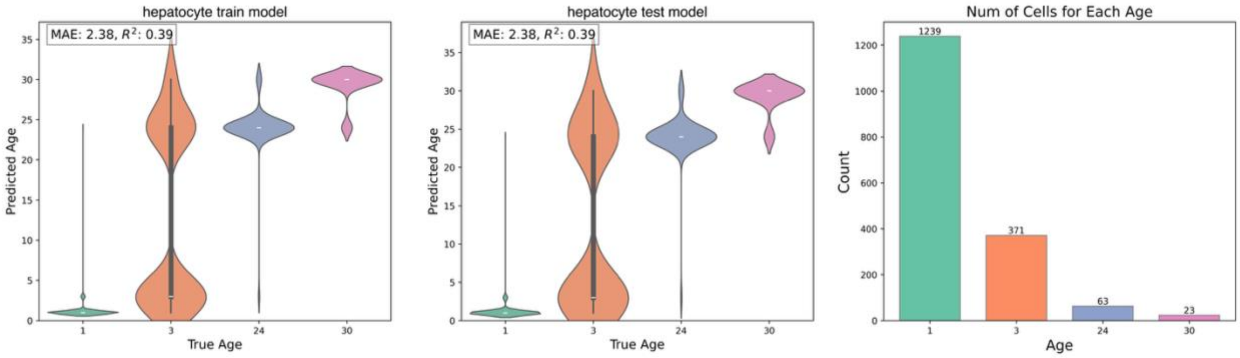


Model using 8192 genes

Liver Kupffer cell Model

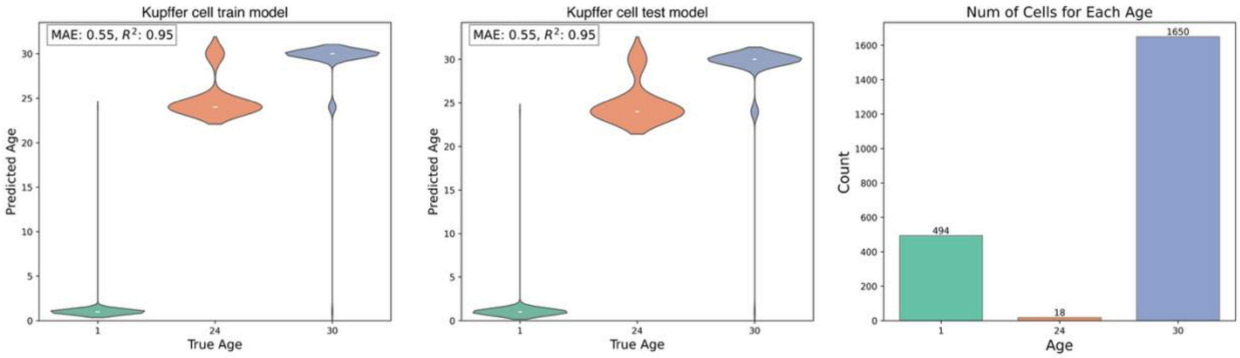


Model using 8192 genes

Lung endothelial cell Model

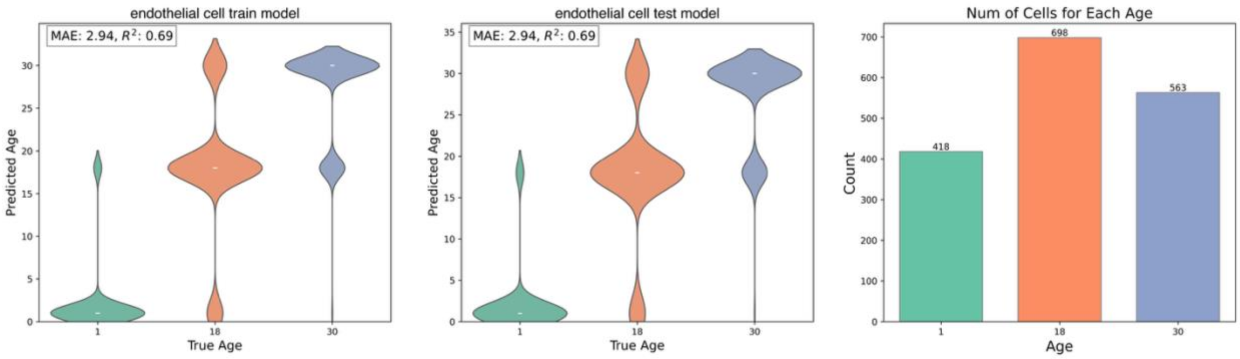
