## Supplementary Figure 9 for "Determining the age of single cells using scMLEAge"

Model using 4096 genes

### Lung lymphocyte Model

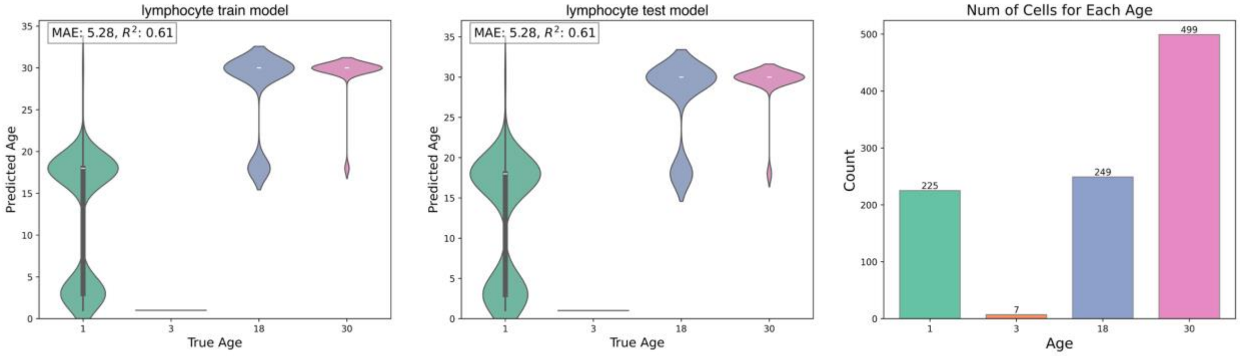


Model using 8192 genes

### Lung macrophage Model

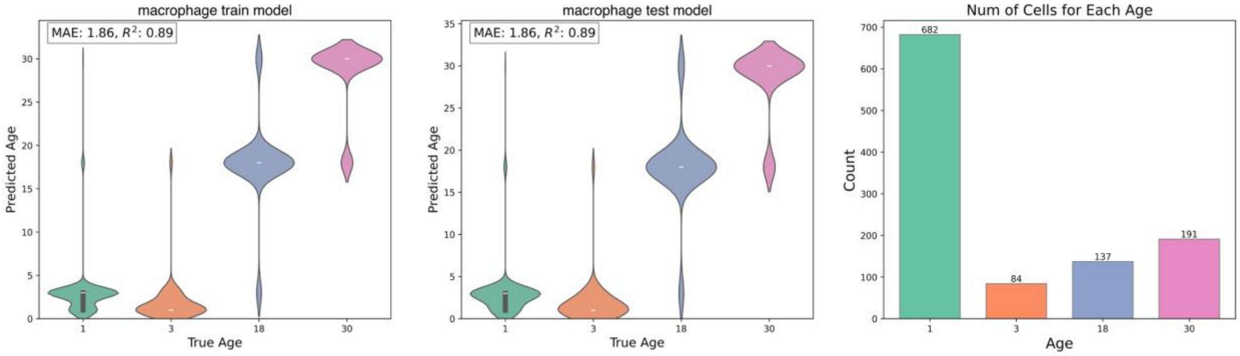


Model using 2048 genes

### Lung monocyte Model

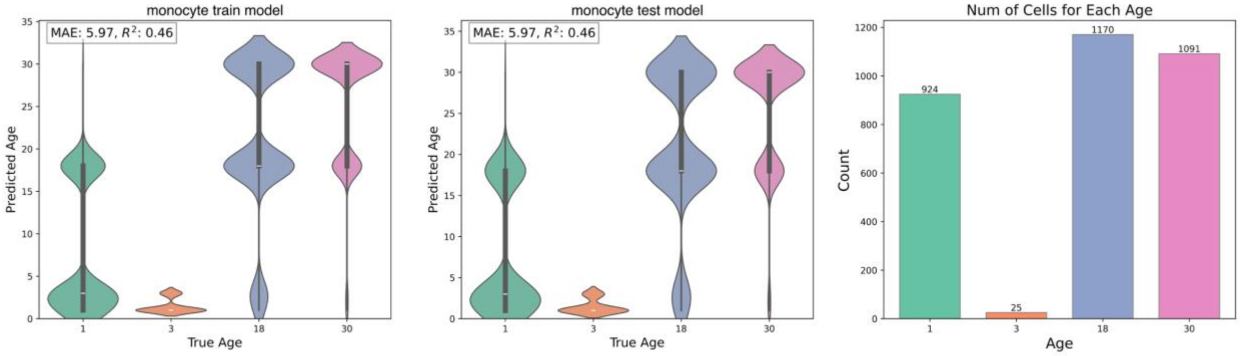


Model using 8192 genes

### Lung stromal cell Model

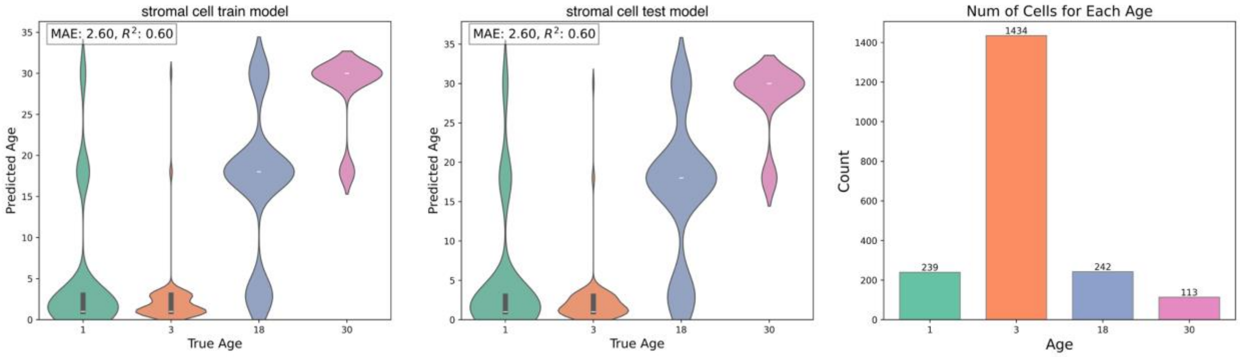
